# Epitope-tagging of the endogenous murine BiP/GRP78/*Hspa5* locus allows direct analysis of the BiP interactome and protein misfolding *in vivo*

**DOI:** 10.1101/2020.01.01.892539

**Authors:** Yunqian Peng, Zhouji Chen, Insook Jang, Peter Arvan, Randal J. Kaufman

## Abstract

BiP/GRP78, encoded by the *Hspa5* gene, is the major HSP70 family member in the endoplasmic reticulum (ER) lumen, and controls ER protein folding. The essential functions of BiP in facilitating proper protein folding are mainly mediated through its dynamic interaction with unfolded or misfolded client proteins, and by serving as a negative regulator of the Unfolded Protein Response. A mechanistic understanding of the dynamics of BiP interaction with its protein partners is essential to understand ER biology, and therefore, we have sought to develop a tractable model to study misfolded protein interaction with BiP. For this purpose, we have used homologous recombination to insert a 3xFLAG epitope tag into the endogenous murine *Hspa*5 gene, just upstream from the essential KDEL signal necessary for ER localization of BiP. Tagging BiP in this way did not alter *Hspa5* expression under basal or ER-stress induced conditions in hepatocytes *ex vivo* or in fibroblasts. Furthermore, the tag did not alter the cellular localization of BiP or its functionality. All of these findings in primary tissue culture were also confirmed *in vivo* in livers of heterozygous mice harboring one WT and one FLAG-tagged *Hspa5* allele. Hepatocyte-specific BiP-FLAG modification did not alter liver function or UPR signaling. Importantly, immunoprecipitation with anti-FLAG antibody completely pulled down FLAG-tagged BiP from lysates of BiP-FLAG expressing livers. Affinity purification-mass spectrometry (AP-MS) of BiP-FLAG protein complexes isolated from the BiP-FLAG-expressing livers of tunicamycin (TM)- and vehicle-treated mice revealed a marked increase in interaction of glycoproteins with BiP-FLAG in response to inhibition of N-glycosylation due to TM-treatment, validating utility of our BiP-Flag mice as a tool to identify ER misfolded proteins *in vivo*. Significantly, our AP-MS analysis also provided *in vivo* evidence demonstrating that BiP-FLAG binds to UPR transducers IRE1α and PERK under basal conditions but is released upon TM-treatment to activate UPR. We have also employed this mouse model to demonstrate that proinsulin in pancreatic β cells misfolds and interacts with BiP-FLAG in healthy mice. In summary, we generated a novel model that can be used to investigate the BiP interactome *in vivo* under physiological and pathophysiological conditions in a cell type-specific manner. This tool provides, for the first time, an unbiased approach to identify unfolded and misfolded BiP-client proteins, and a new approach to study ER protein misfolding in a cell-type and temporal manner to uncover the role of BiP in many essential ER processes.

## INTRODUCTION

Protein misfolding is a protein-specific error-prone process in all cells. In particular, ~30% of all cellular proteins are directed into the endoplasmic reticulum (ER). Proteins enter the ER in an unfolded state and their folding in the ER is challenging because it requires many chaperones and catalysts to assist folding and prevent aggregation in the densely packed unfavorable environment comprised of oxidizing conditions, fluctuating Ca^2+^ concentrations and requiring both proper disulfide bond formation and post-translational modifications(1). Significantly, only proteins that achieve their appropriate 3-dimensional structures can traffic to the Golgi apparatus because of an exquisitely sensitive mechanism that identifies misfolded proteins and retains them in the ER for further productive protein folding or targets them to the degradation machinery mediated by the cytosolic 26S proteasome or macroautophagy. Protein trafficking in the ER is guided by the addition, trimming and modification of asparagine-linked core oligosaccharides in order to engage lectin-based folding machinery to retain misfolded glycoproteins and/or assist in their proper folding (1). Significantly, accumulation of misfolded proteins in the ER initiates adaptive signaling through the unfolded protein response (UPR), a tripartite signal transduction pathway that transmits information regarding the protein folding status in the ER to the nucleus and cytosol to restore ER homeostasis(2, 3). If the UPR cannot resolve protein misfolding, cells may initiate cell death pathways. Stress induced by accumulation of unfolded or misfolded proteins in the ER is a salient feature of differentiated secretory cells and is observed in many human diseases including genetic diseases, cancer, diabetes, obesity, inflammation and neurodegeneration. To elucidate the fundamental etiology of these diseases it is essential to identify which proteins misfold in response to different stimuli, with a future therapeutic goal to learn how to intervene to prevent misfolding and reduce disease pathogenesis.

The characterization of protein misfolding *in vivo* under different physiological conditions is limited due to the absence of conformation-specific antibodies, which are available for some viral glycoproteins, but are mostly lacking for endogenous cellular proteins. In addition, there is a need for an unbiased approach to identify the full spectrum of unfolded and misfolded proteins in the ER, in order to uncover the extent of misfolding of different protein species during disease progression, as well as the impact of different stimuli that can exacerbate ER protein misfolding. The most reliable surrogate for the misfolding of ER client proteins is their interaction with the “Binding Immunoglobulin Protein” known as BiP (encoded by *HSPA5*) which is a heat-shock protein 70 ER chaperone exhibiting a peptide-dependent ATPase activity. BiP was originally characterized as a protein that binds immunoglobulin heavy chains to maintain them in a folding-competent state prior to their heterodimerization with light chains (4). It was also recognized that glucose-deprivation induces a set of genes encoding glucose-regulated proteins, the most abundant being the ER protein GRP78, which is identical to BiP (5). Further studies demonstrated that BiP expression is induced by protein misfolding in the ER through activation of the UPR.

Intriguingly, increased BiP levels feed-back to negatively regulate further UPR activation. One hypothesis posits that BiP binding to the UPR sensors IRE1, ATF6 and PERK inhibits their signaling (6), although there is no direct evidence to support this notion in a physiological setting *in vivo*. Early studies to analyze protein misfolding demonstrated that only misfolded proteins that bind BiP activate the UPR and those that do not bind BiP do not activate the UPR (7–13). Unfortunately, however, there are no BiP antibodies currently available that can efficiently recognize BiP-client protein complexes in the absence of chemical crosslinkers, thus limiting the ability to study protein misfolding in the ER. As BiP provides many essential ER functions (including regulating Sec61 for co-translational and post-translational translocation into the ER, protein folding and degradation, maintenance of ER Ca^2+^ stores, repressing UPR signaling, *etc.),* characterizing BiP interactions *in vivo* is essential to understand all these processes, and will provide significant insight into the role of ER protein misfolding in disease pathogenesis.

BiP immunoprecipitation (IP) from whole tissue lysates has the limitation that BiP is ubiquitously expressed; thus, IP recovers BiP and its partner proteins from multiple cell types. With this in mind, we have sought the ability to detect cell type-specific BiP interactions at different stages of disease progression. In addition, we wanted to avoid BiP overexpression, because this increases non-physiological BiP interactions (10). Therefore, we used homologous recombination to generate a conditional allele in mice with insertion of a 3xFLAG tag into the C- terminus of the endogenous BiP (*Hspa5*) coding sequence, just upstream from the KDEL ER localization signal. The engineered allele is designed such that upon cell type-specific *Cre*-induced deletion, expression of BiP-3xFLAG from the endogenous locus will permit endogenous BiP expression with the ability to identify BiP-interactors by anti-FLAG IP.

## MATERIALS AND METHODS

### Generation of BiP-3xFlag mice

We generated a conditional knock-in mouse model by modifying of the *Hspa5* locus. This was achieved by floxing a targeted WT exon 9-pA cassette upstream of the knock-in exon 9 where a 3xFLAG sequence was introduced immediately prior to the KDEL ER retention signal. Additionally, a FRT-flanked neomycin cassette was introduced into the floxed region. The genetic modification was introduced into Bruce4 C57BL/6J ES cells (14) via gene targeting. Correctly targeted ES cell clones were identified and then injected into goGermline blastocysts (15, 16). Male goGermline mice were bred to C57BL/6J females to establish heterozygous germline offspring on a C57BL/6J background. **a). Vector construction.** A replacement vector targeting *Hspa5* exon 9 *coding sequence* region (CDS region) was generated by assembly of 4(ABCD) fragments using sequential cloning. The first fragment which encompassed the 3 kb 5’-homology arm was generated by PCR amplified from C57BL/6J genomic DNA using primers P2093_41 and P2093_51. The second and third fragments which comprise loxP-exon9-BGHpA and exon 9-3xFlag were synthesized by Genewiz, respectively. The fourth fragment comprising the 3.2kb 3’-homology arm was generated by PCR amplification from C57BL/6J genomic DNA using primers P2093_44 and P2093_54. Synthesized fragments and PCR primers used to amplify the fragments included all the restriction enzyme sites required to join them together and to ligate them into the Surf2 vector backbone (Ozgene). The final targeting vector 2093_Teak_ABCD contained a FRT-flanked neomycin selection (neo) cassette, an exon 9 *coding sequence* sequentially with an inserted bovine growth hormone (BGH) polyA tail, an additional exon 9 coding sequence sequentially with a 3xFLAG tag cassette right before the KEDL sequence, 5’ and 3’-loxP site (Figure 1A). For sequence information of the primers see Suppl Table 1. The targeting vector was entirely sequenced and then linearized by digestion with PmeI before electroporation into C57BL/6J Bruce4 ES cells (14). Neo-resistant ES cell clones were screened by qPCR to identify potentially targeted clones. **b). Targeting murine ES cells through homologous recombination.** TaqMan® copy number reference assays were used to measure copy number in the genome. Two pairs of primers were used to amplify the WT locus at the extreme 5’ and 3’ positions to detect 2 copies from the WT allele and 1 copy from the targeted allele (primers, 2093_Lo5WT and 2093_LoWT3). Another primer pair targeting Neo sequence was used to test the targeted allele (primer, 1638_goNoz). Two genes from Y chromosome (1 copy) and chromosome 8 (2 copy) were used as control. Two positive clones, Clones I_1D08 and I_1G08, were confirmed as correctly targeted and were used to injection into goGermline blastocysts**. c). Production of mice heterozygous for a BiP-FLAG allele.** ES cells from clones I_1D08 and I_1G08 were injected into goGermline donor blastocysts to generate chimeras. A total of 84 injected blastocysts were transferred into 7 recipient hosts. These resulted in 35 offspring, of which 28 were male chimeras. Four males were chosen for mating with homozygous *Flp* mice. A total of 17 pups was born from three litters, including 10 WT and 7 WT/conKI (Figure 1B).

**Figure 1.**
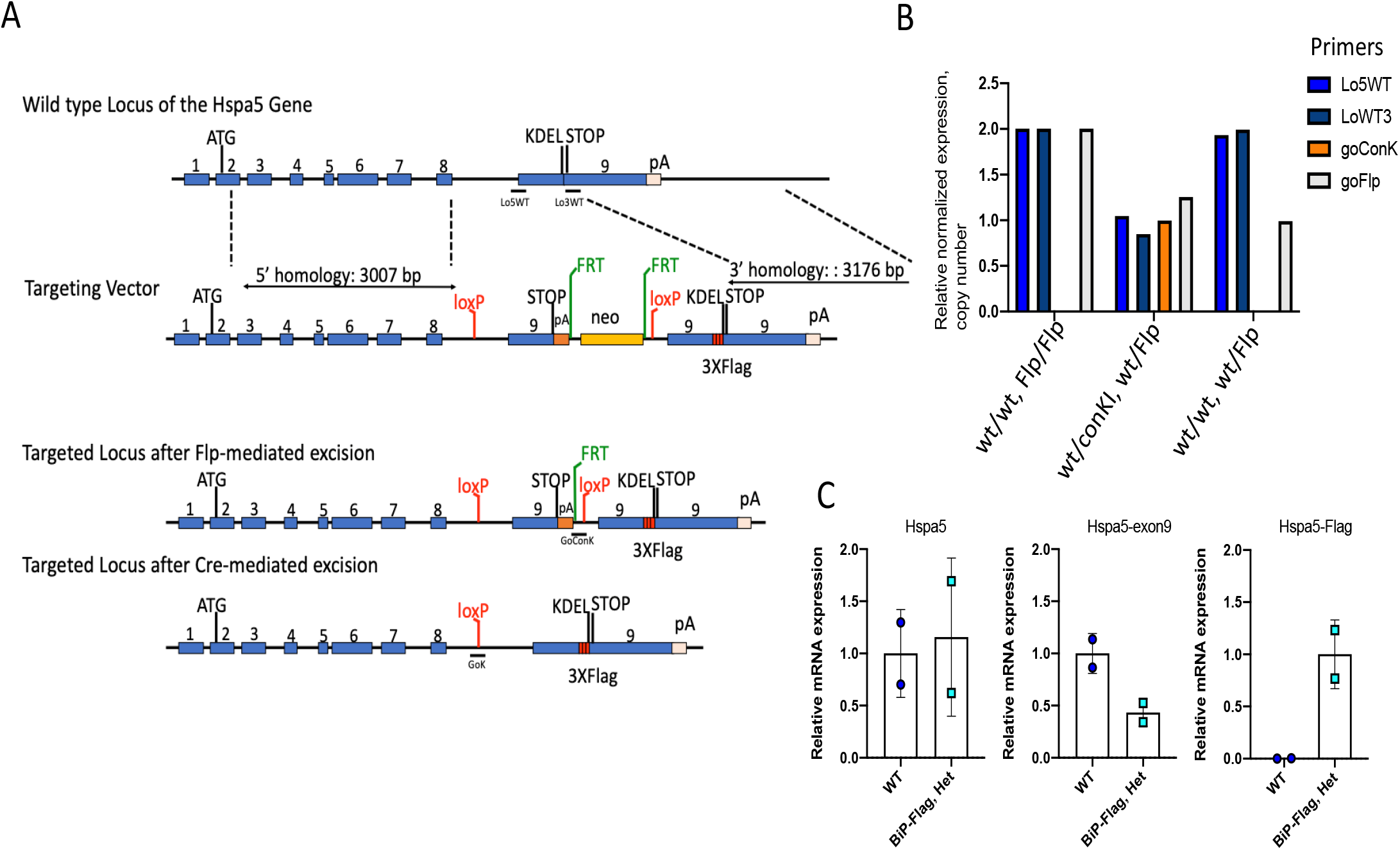
BiP-FLAG-Het mice were generated by homologous recombination. **A. Targeting strategy.** The replacement vector contains a Neo-cassette (yellow) flanked by FRT sites (green) which, together with targeted WT exon 9-pA cassette, are flanked by LoxP sites (red) followed by duplicated exon 9 containing a 3xFLAG sequence(red) upstream from the KDEL motif. Using 5’ and 3’ homology arms the vector was used to target the Hspa5 locus of murine ES cells. The Neo-cassette was removed by Flp-mediated recombination. After Cre mediated recombination, exon 9-FLAG is expressed under control of the endogenous Hspa5 regulatory elements. Blue boxes denote exons. Lo5WT, Lo3WT and GoConK represent qPCR probes for the specified locus. **B. Copy number quantification.** qPCR was performed to analyze BiP-FLAG-Het mice. qPCR reactions display genotyping from WT/WT-Flp/Flp, WT/conKI-WT/Flp (mice number, 2093_044_A048F), and WT/conKI-WT/Flp(mice number, 2093_044_A047F) mice. Primers are indicated in panel A. Lo5WT and LoWT3 identify the WT allele of Hspa5. goConK identifies the knock-in allele of Hspa5. **C. Targeted allele expression.** Total RNA was extracted from BiP-FLAG-Het mice liver infected with AAV8-TBG-Cre. mRNA expression of Hspa5 was measured by qRT-PCR.

### Isolation and culture of primary hepatocytes and skin fibroblasts

Mouse primary hepatocytes were isolated by portal vein perfusion of collagenase as described (17). Murine skin fibroblasts were prepared by collagenase (Type II and Type IV, Sigma) digestion of abdominal skins dissected from a female BiP-FLAG-Heterozygous (Het) mouse and an *Hspa5* wild type littermate (6-weeks old). The primary hepatocytes and skin fibroblasts were cultured in DMEM/10% FBS. After overnight culture, cells were transduced with Ad-βGal or Ad-Cre at an MOI of 34. Where specified, cells were treated with castanospermine (CST, 20 μM) or tunicamycin (Tm, 0.5 μg/ml) for 22 h to induce ER stress.

### Mouse experiments

Four female BiP-FLAG-Het mice and 4 of their female littermates were used for an *in vivo* experiment. They were infused with AAV8-TBG-Cre (2.5 × 10^11 vg/mouse) through tail vein injection at 6.5 wks of age. After 10 days, mice were treated with Tm (1 mg/Kg) or vehicle (saline) through I.P. injection and were sacrificed for tissue collection after 17 h.

### qRT-PCR and qPCR analyses

Total RNAs were extracted from isolated liver by RNeasy Mini Kit (Qiagen). cDNAs were synthesized by iScript cDNA Synthesis kit (Bio-Rad Laboratories, Inc). The relative mRNA levels were measured by qRT-PCR with iTaq Universal SYBR green Supermix (Bio-Rad Laboratories, Inc). All primers are listed in Suppl. Table 1.

### Immunofluorescence microscopy

Cells were plated on coverslips for overnight and fixed with 4%PFA. Cells and Sections were stained with the following antibodies; FLAG (M2, Sigma), α-PDIA6 (18233-1-AP, Proteintech), and DAPI (Fisher Scientific). For secondary antibodies we used: Alexa Fluor 488 goat α-rabbit IgG, Alexa Fluor 594 goat α-mouse IgG, anti-bodies (Invitrogen). Images were taken by a Zeiss LSM 710 confocal microscope with a 20X and 63X objective lenses. Scale bars are indicated in the figures.

### Western blot analyses

All Western blots were performed separating proteins by SDS-PAGE on a 5-15% gradient polyacrylamide gel for transfer onto nitrocellulose membranes, followed by blocking with Licor Blocking solution and incubation with primary and fluorescent-labeled secondary antibodies (Licor). The immune-fluorescence signals were captured using a Licor scanner. The key primary antibodies used in this study were as follows: Flag (M2, Sigma), BiP(3177, CST), KDEL (SC-58774, SCBT), PDIA4 (14712-1-AP, Proteintech), PDIA6 (18233-1-AP, Proteintech), β-Actin (8H10D10, CST).

### Affinity purification-mass spectrometry (AP-MS) of BiP-FLAG protein complexes

Liver samples were lysed in a buffer containing 50 mM Hepes-NaOH pH 8.0, 100 mM KCl, 2 mM EDTA, 0.5% NP40 and 10% glycerol, supplemented with protease and phosphatase inhibitor cocktails. The liver lysates were centrifuged at 15,000 g for 20 min at 4 C. The resultant supernatants were subjected to IP with M2 anti-FLAG magnetic beads (Sigma). A sample of beads was removed for Western blotting and the majority of beads were subjected to denaturation, reduction and overnight trypsin/lys-C mix digestion for 2D LC-MS/MS (18). MS/MS spectra were searched against the Mus musculus Uniprot protein sequence database using Maxquant (version 1.5.5.1) with false discovery rate (FDR) set to 1%. MSStats was used to calculate a confidence (p-value) and fold change of BiP-FLAG Het_Tm IP/BiP-FLAG Het_Veh after correction with Wt_Tm IP and Wt_Veh IP, respectively.

### Glucose tolerance tests

Mice were fasted for 5 hr before IP injection of glucose (2 g/Kg body weight). Blood glucose levels were measured by tail-bleeding at 0, 15, 30, 45, 60, and 90 min.

### Pancreas immunohistochemistry

Pancreata were harvested and fixed in 4% PFA. Paraffin embedding, sectioning, and slide preparations were performed in the SBP Histopathology Core Facility. Sections were stained with primary antibodies: Flag (M2, Sigma), PDIA6 (18233–1-AP, Proteintech), Insulin (Guinea pig polyclonal anti-Insulin antibody was produced in-house) and secondary antibodies: Alexa Fluor 594 goat α-guinea pig IgG, Alexa Fluor 488 goat α-mouse IgG, Alexa Fluor 594 goat α-mouse IgG, Alexa Fluor 488 goat α-rabbit IgG (Invitrogen). Images were obtained by Zeiss LSM 710 confocal microscope with a 40X objective lens. Scale bar, 20 μm.

### Islet Isolation

Islets were isolated as previously described(19). Islets were handpicked and analyzed directly or after overnight recovery in RPMI 1640 media (Corning 10-040-CV) supplemented with 10% FBS, 10 mM Hepes, 1mM Sodium pyruvate, 1% penicillin/streptomycin, and 100μg/ml primocin.

### Islet Western blotting

Isolated islets were briefly rinsed in PBS and lysed in RIPA buffer (10 mM Tris pH 7.4, 150 mM NaCl, 0.1% SDS, 1% NP-40, 2 mM EDTA) with protease and phosphatase inhibitors on ice for 10 min. Lysates were obtained by centrifugation at 12000 g for 10 min at 4°C. Samples were prepared in Laemmli sample buffer with (reducing) or without (non-reducing) 5% β-mercaptoethanol, boiled at 95°C for 5 min, and analyzed by SDS-PAGE (4-12% Bis-Tris gel, Bio-Rad Laboratories, Inc). Non-reducing gels were incubated in 10 mM DTT for 5 min at room temperature prior to transfer to nitrocellulose membranes. Primary antibodies used in this study: Vinculin (V9131, Sigma), Flag (M2, Sigma), BiP (3177, Cell Signaling Technology), BiP & GRP170 (friendly gift from Dr. Hendershot), PDIA4 (14712–1-AP, Proteintech), PDIA6 (18233-1-AP, Proteintech), p58IPK (2940S, Cell Signaling Technology), ERdj3 (13157-1-AP, Proteintech), proinsulin (HyTest Ltd., 2PR8, CCI-17). Secondary antibodies were used in 1:5000 (Li-Cor, IRDye-800CW or IRDye-680RD).

### Immunoprecipitation of islet lysates

Isolated islets were briefly rinsed in PBS containing 20 mM N-ethylmaleimide (NEM) and lysed in HG lysis buffer (20 mM Hepes-KOH pH 7.4, 150 mM NaCl, 2 mM MgCl2, 10 mM D-Glucose, 10% glycerol, 1% Triton-X-100) with 100 U/ml Hexokinase,2 mM NEM, and protease and phosphatase inhibitors on ice for 10 min. Lysates were obtained by centrifugation at 12000 g for 10 min at 4°C and incubated with goat-anti-Flag agarose beads (PA1-32374, Invitrogen) or proinsulin Ab (CCI-17) bound protein G beads (GE17-0618-01, Sigma) for overnight at 4°C. Beads were eluted in 2X Laemmli sample buffer with (reducing) or without (non-reducing) 5% β-mercaptoethanol. IP elutes were analyzed by SDS-PAGE according to islet Western blotting methods.

## RESULTS

### Generation of BiP-FLAG mice

BiP-FLAG conditional knock-in mice were generated by targeting exon 9 region and flanking of with LoxP sites via gene targeting in Bruce4 C57BL/6J embryonic stem (ES) cells (14). Gene-targeted ES cell clones were identified, and cells then injected into goGermline blastocysts (15, 16). Male chimeric mice were bred with Flp female mice to delete the Neo cassette and establish heterozygous germline offspring on a C57BL/6J background (Figure 1A). TaqMan® copy number assay was used to genotype the offspring (Figure 1B). A total of 17 pups was born from three litters, including 10 WT and 7 WT/conKI (41% observed vs. 50% expected). All of these pups grew normally and appeared healthy. No difference in body weights between genotypes was observed (Suppl. Figure 1).

To test whether protein expressed from the Cre-induced *Hspa5-FLAG* allele is similar to the endogenous *Hspa5* allele, we delivered AAV-Cre by intravenous tail vein injection into mice as described for the *in vivo* experiments. mRNA expression from the endogenous and targeted *Hspa5* alleles was measured by qRT-PCR with primers directed at the targeted region, including crossing the FLAG insertion. With AAV-Cre induced LoxP deletion in liver, WT mice demonstrated an ~2-fold increased expression compared to the BiP-FLAG-Het mice with primer *Hspa5-exon 9* that identifies the WT allele, as expected. While the other *Hspa5* primer that does not target the FLAG insertion did not show significant difference between the WT and knock-in mice. The primer targeting the FLAG sequence was only observed upon amplification in BiP-FLAG-Het mice. This confirmed the FLAG knock-in into the *Hspa5* locus at exon 9 (Figure 1C).

### BiP-FLAG expression *ex vivo* in primary hepatocytes and skin fibroblasts isolated from BiP-FLAG heterozygous (Het) mice

To activate BiP-FLAG expression *ex vivo*, we isolated primary hepatocytes and skin fibroblasts from heterozygous BiP-FLAG mice and transduced them with Ad-Cre to induce Cre-mediated deletion of the floxed *Hspa5* segment. At 24 h after Ad-Cre-transduction, approximately 90% of the BiP-FLAG-Het hepatocytes and fibroblasts were positive for FLAG immunofluorescence (Suppl. Figure 2).

In hepatocytes, Western blot analysis detected BiP-FLAG migrating slightly above endogenous BiP in the BiP-FLAG-Het hepatocytes as early as 22 h after Ad-Cre transduction (not shown). By 3 days after Ad-Cre transduction, the steady-state-level of BiP-FLAG was about 50% of the endogenous BiP under basal conditions but increased to a level similar to that of the endogenous BiP produced from the untargeted *Hspa5* allele after Tm-treatment for 20 h, based on Western blotting analysis with a rabbit anti-BiP monoclonal antibody (Figure 2), suggesting that the 3xFLAG insertion into the BiP C-terminus does not alter *Hspa5* expression in our mouse model. Significantly, very similar levels of total BiP were observed in the Ad-βGal- and Ad-Cre-transduced BiP-FLAG-Het hepatocytes at 20 h after Tm-treatment (Figure 2), suggesting that our genetic modification of *Hspa5* did not alter UPR signaling for BiP induction.

**Figure 2.**
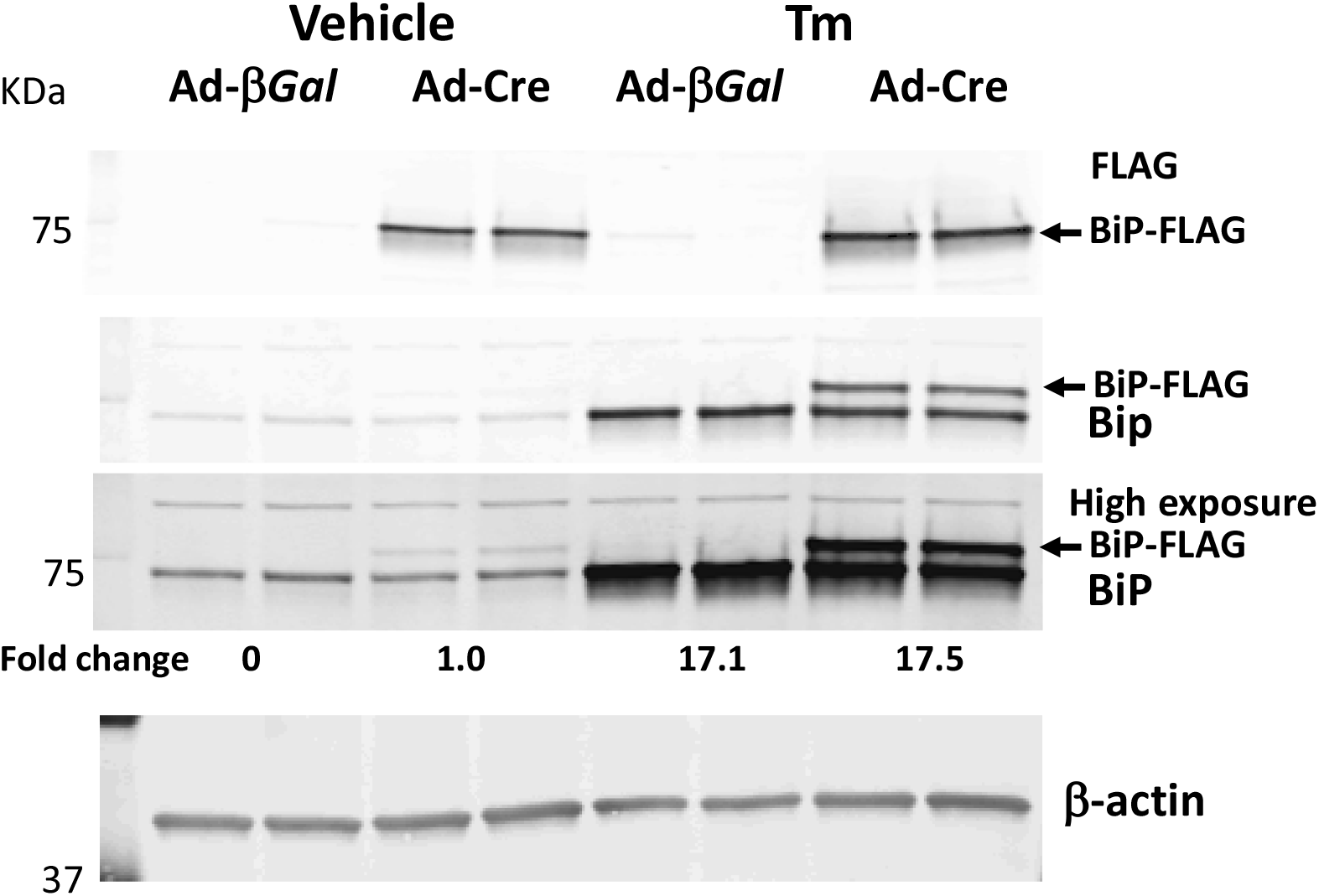
The BiP-FLAG allele is induced by ER stress in primary hepatocytes. Hepatocytes were isolated from a 6-wk old female BiP-FLAG-Het mouse, plated onto 24 well plates and infected with the indicated adenoviruses at 4 h after plating. After 48 h, cells were treated with Tm 0.5μg/ml or vehicle for 20 h and then harvested for Western blot analysis.

Unlike primary hepatocytes, Ad-Cre activation of the BiP-FLAG knock-in locus in BiP-FLAG-Het primary skin fibroblasts resulted in equal levels of endogenous BiP and BiP-FLAG (Figure 3). This difference between skin fibroblasts and primary hepatocytes may be explained by the fact that fibroblasts, but not hepatocytes, proliferate *in vitro*, leading to a dilution of the preexisting endogenous BiP in fibroblasts. Significantly, the increase in endogenous BiP and BiP-FLAG were nearly identical in response to Tm or castanospermine (CST), to inhibit α- and β-glucosidases, to activate the UPR (Figure 4).

**Figure 3.**
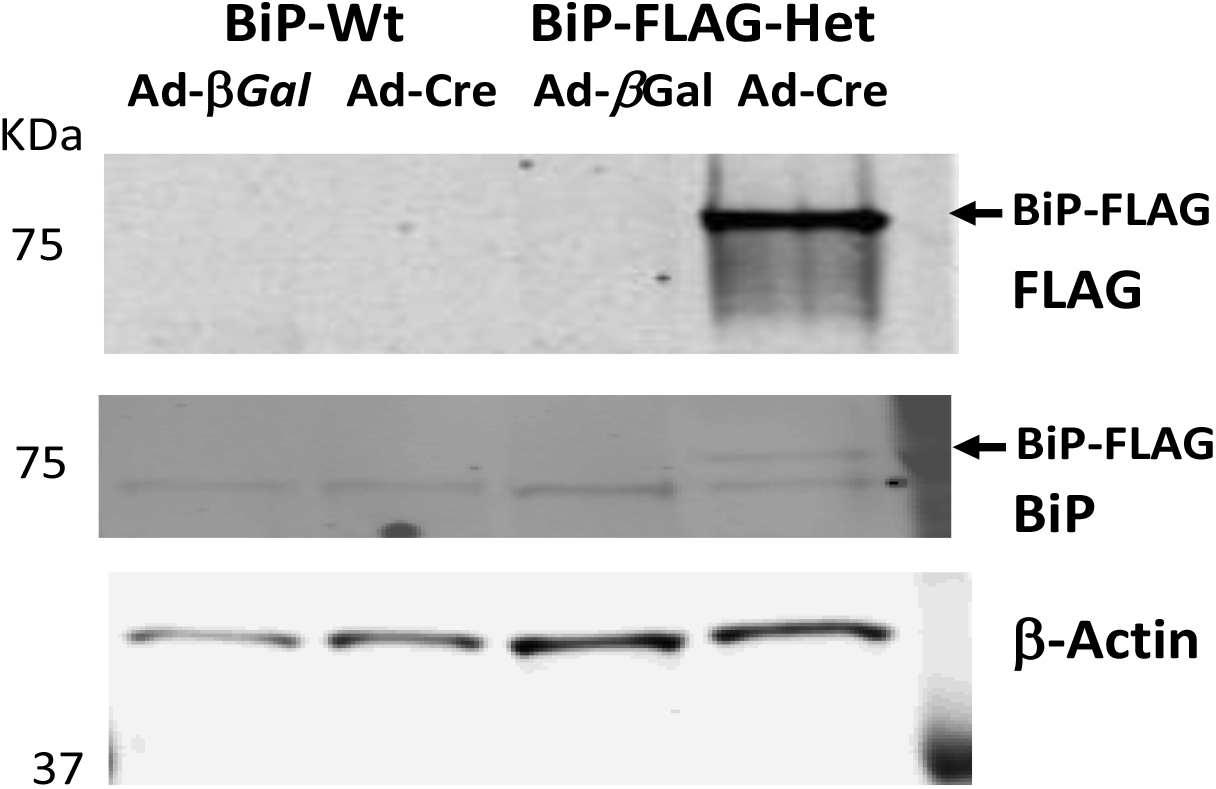
BiP-FLAG is activated by Ad-Cre infection in BiP-FLAG primary fibroblasts. Skin fibroblasts were isolated from a 6-wk old female BiP-FLAG-Het mouse and plated onto 24 well plates, infected with the indicated adenoviruses after the first passage and harvested for Western blot analysis.

**Figure 4.**
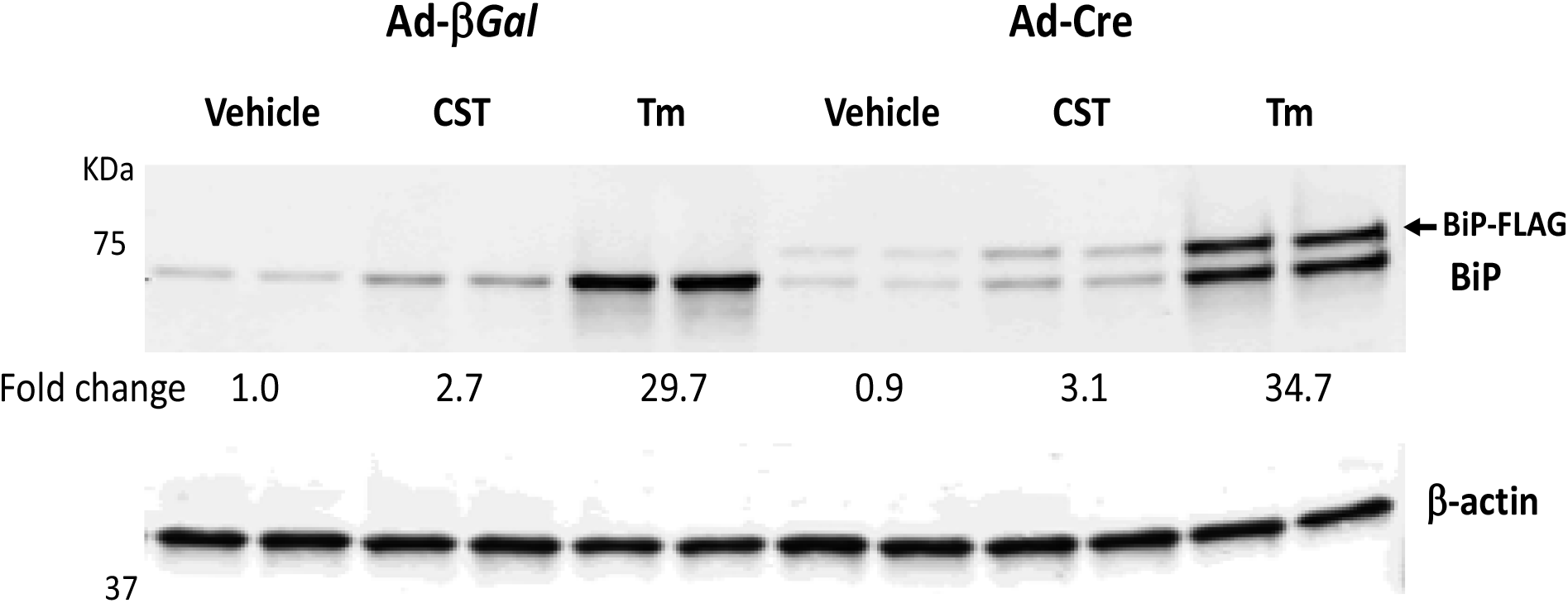
BiP-FLAG is induced in BiP-FLAG primary fibroblasts in response to ER stress. Skin fibroblasts were isolated from a 6-wk old female BiP-FLAG-Het mouse, plated onto 24 well plates, infected with the indicated adenoviruses after the first passage. After Ad-infection, cells were treated with Tm 0.5 μg/ml, CST 20 μM or vehicle for 22 h and harvested for Western blot analysis.

### BiP-FLAG is localized to the ER

An essential question is whether tagging the C-terminus of BiP may alter its intracellular localization as the FLAG tag is adjacent to the KDEL ER retention signal. Immunofluorescence microscopy showed that BiP-FLAG colocalized with the ER localized PDIA6 both in Ad-Cre-transduced BiP-FLAG-Het primary hepatocytes (Figure 5A) and skin fibroblasts (Figure 5B), importantly demonstrating that insertion of the 3xFLAG tag into BiP did not alter its cellular localization.

**Figure 5.**
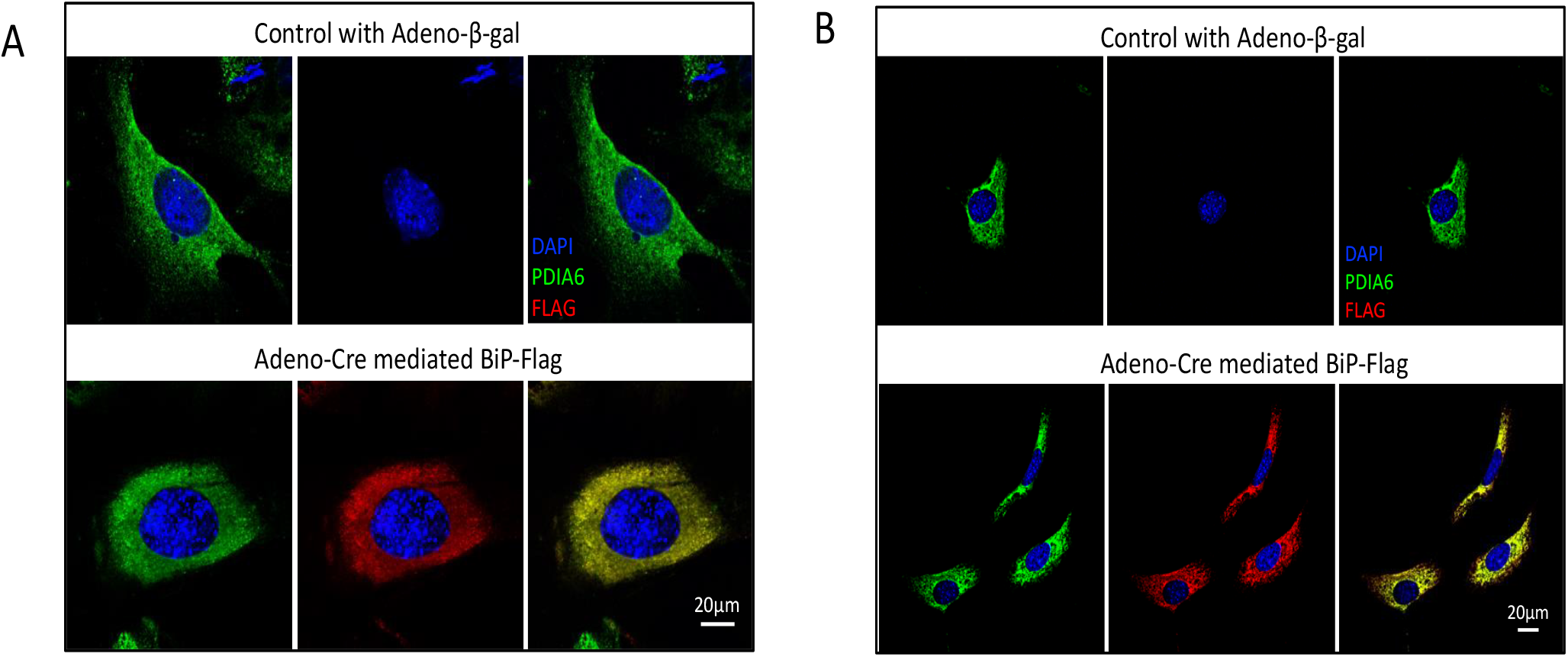
BiP-FLAG is localized to the ER lumen. **A.** Hepatocytes were isolated from a 6-wk old female BiP-FLAG-Het mouse, plated onto 6 well plates and infected with the indicated adenoviruses at 4 h after plating. After 4 days, cells were fixed with formalin and stained with anti-FLAG antibody, anti-PDIA6 antibody and DAPI. Images were captured by a 63 oil lens from each group by confocal microscopy. Scale bar, 20 μm. **B.** Fibroblasts were isolated from a 6-wk old female BiP-FLAG-Het mouse, plated onto 6 well plates and infected with the indicated adenoviruses at 7 days after plating. At 24 h after Ad-infection, cells were fixed with formalin, stained with antibodies for FLAG or PDIA6 and for DAPI. Images were captured by a 63 oil lens from each group by confocal microscopy. Scale bar, 20 μm.

### Hepatocyte-specific Cre-expression in BiP-FLAG-Het mice demonstrates intact functional activities of BiP-FLAG *in vivo*

Together, the above findings show that BiP-FLAG knock-in did not alter the expression, localization or the functional activity of the endogenous or the modified *Hspa5* alleles. To confirm these findings *in vivo* and to explore the feasibility for hepatocyte-specific BiP-FLAG knock-in, we infused AAV8-TBG-Cre into 4 BiP-FLAG-Het mice and 4 WT littermates to express Cre selectively in hepatocytes. The TBG promoter is a hybrid promoter comprised of the human thyroxine-binding globulin promoter and microglobin/bikunin enhancer that is specifically expressed in hepatocytes. These mice were treated with Tm (1 mg/Kg) or vehicle (saline) at day 10 after AAV8-infusion for 17 h. Cre-mediated activation of BiP-FLAG in hepatocytes of BiP-FLAG-Het mice did not alter plasma or hepatic lipid levels and did not alter liver morphology in the absence or presence of Tm-treatment (Figure 6A–C, Suppl. Figure 3). In addition, BiP-FLAG was detected in nearly all hepatocytes in the livers of both AAV8-TBG-Cre-infused BiP-FLAG-Het mice.

**Figure 6.**
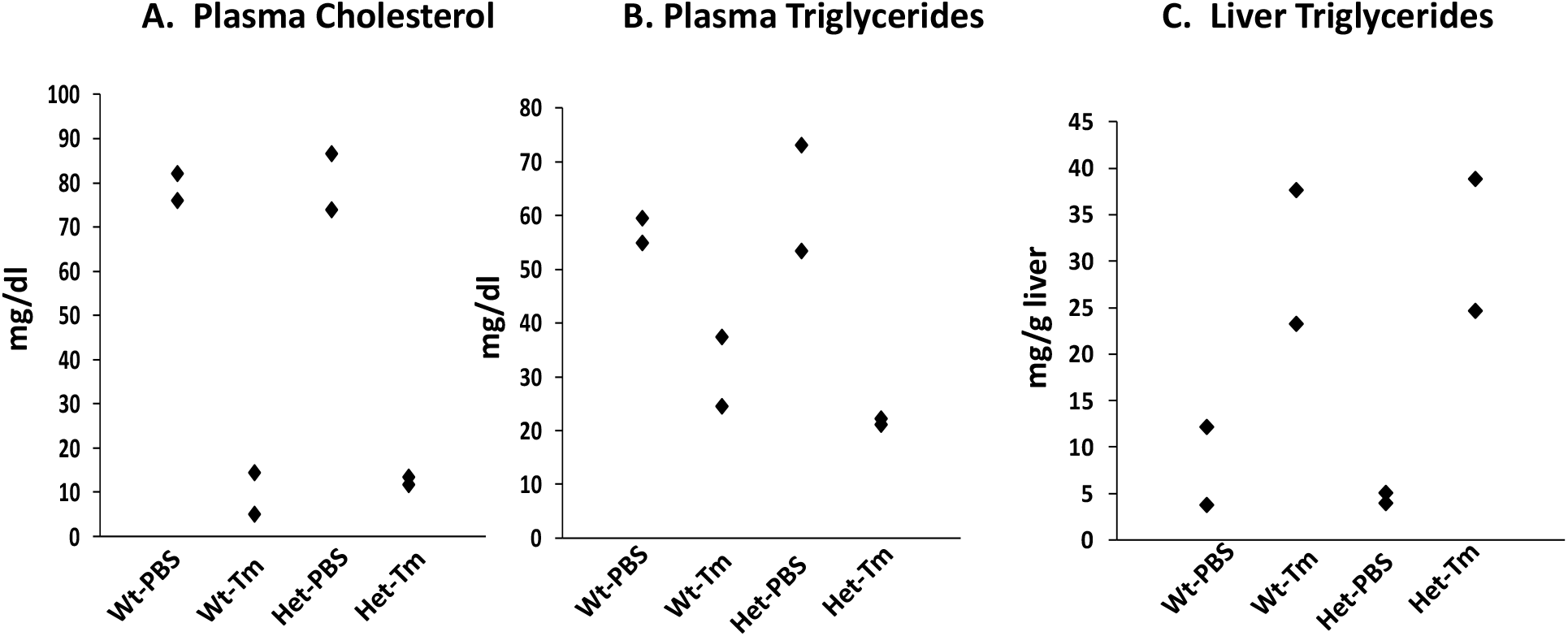
AAV8-TBG-Cre-treatment of BiP-FLAG-Het mice does not alter plasma or hepatic lipid contents. Plasma cholesterol and triglyceride levels and hepatic triglyceride contents of AAV8-TBG-Cre-treated *Hspa5* wildtype (WT) and BiP-FLAG-Het (Het) mice at 17 h after injection with vehicle (PBS) or Tm were measured. Each data point represents one individual mouse.

Like the Ad-Cre-transduced BiP-FLAG-Het skin fibroblasts, there were equivalent levels endogenous BiP and BiP-FLAG in the livers of the AAV8-TBG-Cre-treated BiP-FLAG-Het mice with or without Tm-treatment (Figure 7). Importantly, BiP-FLAG knock-in did not alter expression of GRP94, PDIA4 and PDIA6 as well as BiP in the livers under basal or Tm-induced conditions (Figure 7). qRT-PCR assay demonstrated that activation of the *Hspa5*-FLAG allele did not alter expression of key UPR genes under basal or induced conditions (Figure 8).

**Figure 7.**
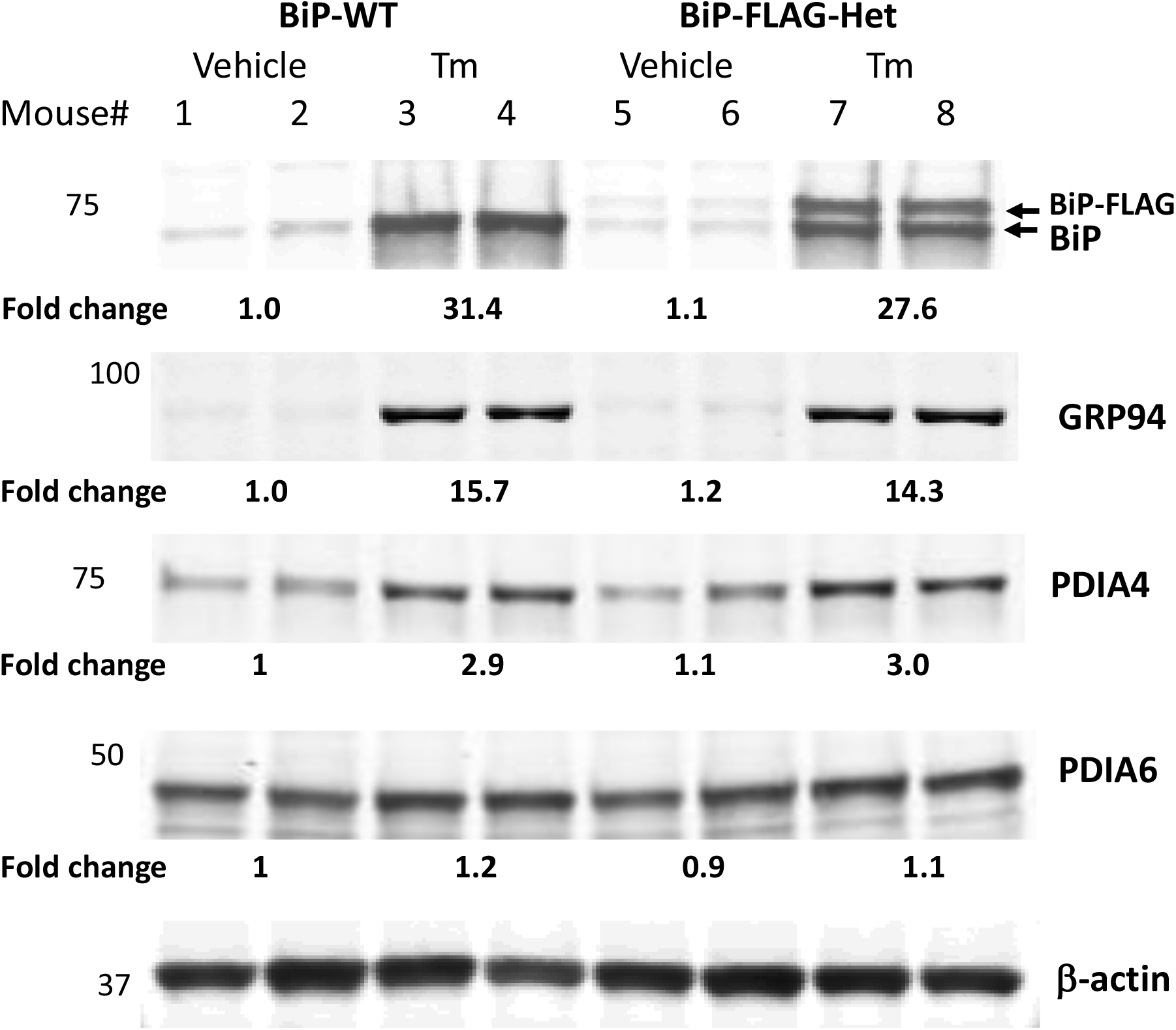
BiP-FLAG induction in hepatocytes by ER stress *in vivo.* After infection with AAV8-TBG-Cre or control virus for 10 days, mice were treated with Tm (1 mg/Kg) or vehicle (saline). After 17 h, liver tissues were collected immediately after sacrifice and for lysis in RIPA buffer. Each lane in the Western blot represents an individual mouse.

**Figure 8.**
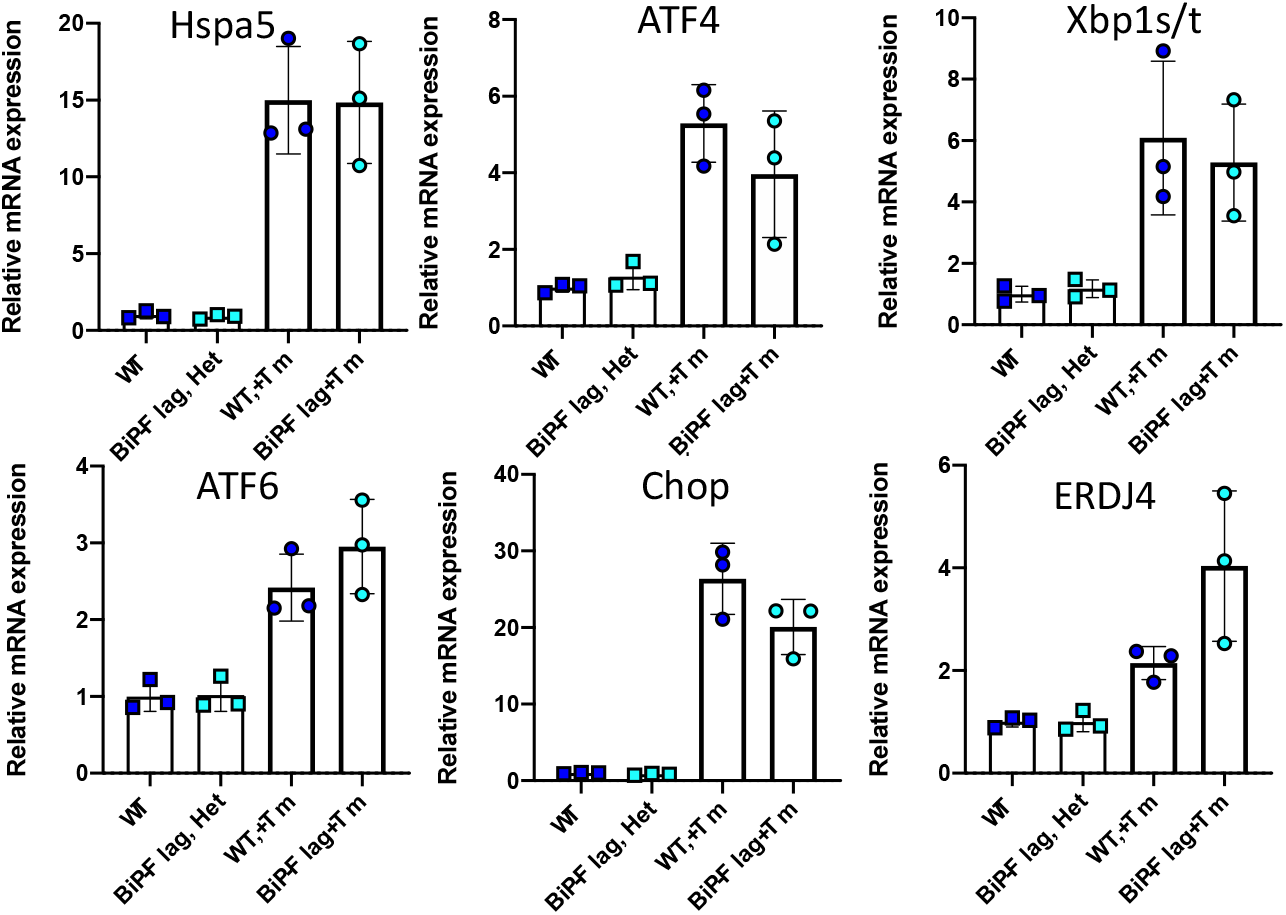
Hepatocyte-specific BiP-FLAG knock-in does not alter expression of key UPR genes in the liver. Total RNAs were isolated from liver samples collected as described in Figure 7 and subjected to qRT-PCR analysis to measure mRNA levels for the indicated genes normalized to 18S rRNA.

Finally, to confirm the ability of anti-FLAG antibody to immunoprecipitate (IP) BiP-FLAG synthesized *in vivo*, we performed FLAG-IP assays of liver lysates prepared from the BiP-FLAG-Het mice. A mouse anti-FLAG antibody completely depleted BiP-FLAG from the IP supernatants of the AAV8-TBG-Cre-treated BiP-FLAG-Het liver lysates (Figure 9), demonstrating a high efficiency for BiP-FLAG pulldown. While more work is necessary to understand the nature and composition of the proteins pulled down with BiP-FLAG, the finding that a significant amount of endogenous BiP was pulled down with BiP-FLAG from the liver lysates of the AAV8-TBG-infected BiP-FLAG-Het mice, especially those with Tm-treatment (Figure 9), indicates that BiP-FLAG was pulled down as protein complexes.

**Figure 9.**
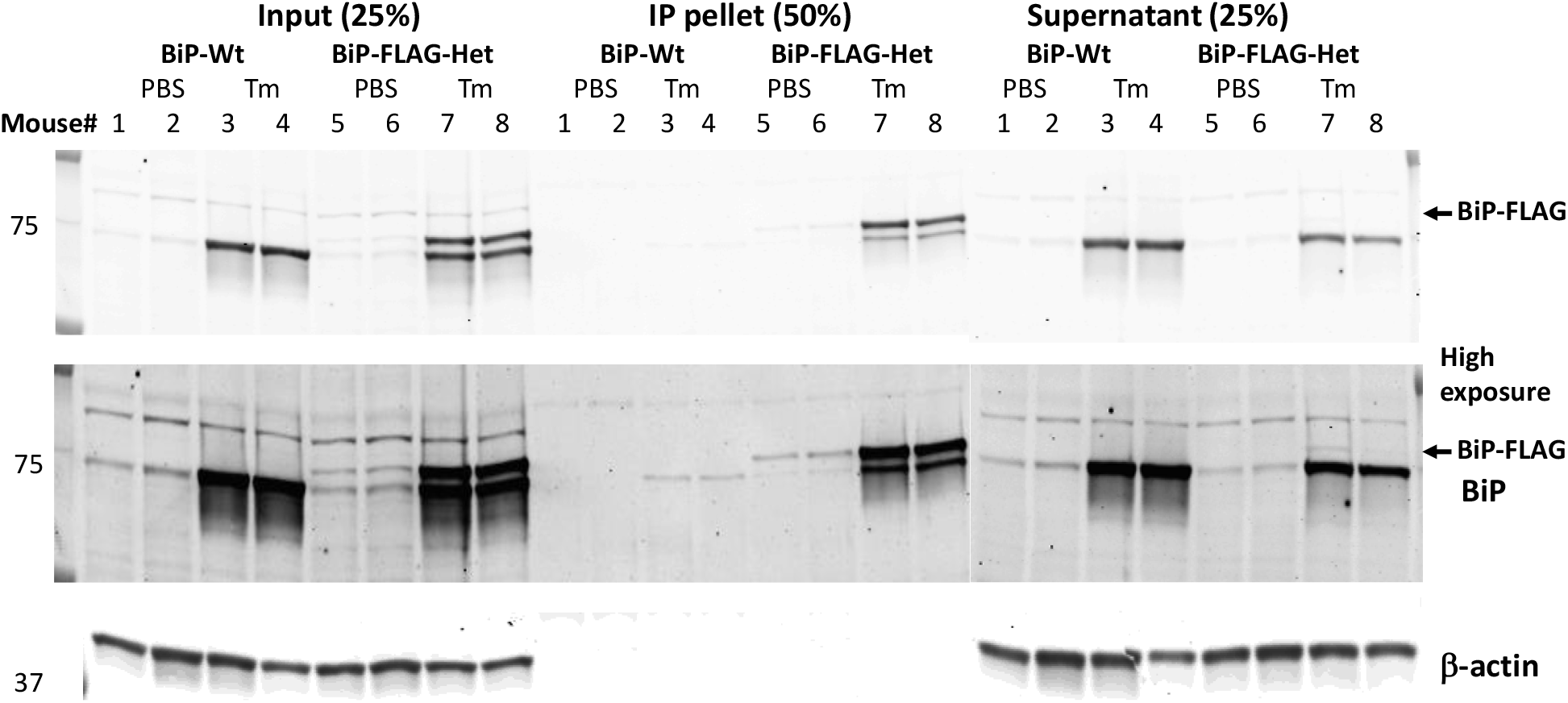
BiP-FLAG is efficiently pulled down from AAV8-TBG-Cre/BiP-FLAG-Het liver lysates using murine anti-Flag (M2) agarose. Liver lysates prepared as described in Figure 7 were subjected to IP using anti-FLAG (m2)-coupled to agarose. Western blotting was performed using a rabbit anti-BiP monoclonal antibody (3177, CST) as a primary antibody.

### Identification of misfolded proteins in hepatocytes in a murine model of ER stress

To characterize the protein complexes that were pulled down with BiP-FLAG from lysates of the BiP-FLAG-expressing murine livers through FLAG-affinity purification, we performed affinity purification-mass spectrometry (AP-MS) analysis on livers of Tm- or vehicle-treated BiP-FLAG heterozygous mice, with Tm- or vehicle-treated wild type littermates as negative controls (Figure 9).

Figure 10A shows all proteins that had increased interaction with BiP-FLAG in response to Tm treatment. Importantly, most of these proteins are N-glycosylated proteins, *e.g.,* insulin receptor (*Insr*) and EGF receptor (*Egfr*) in the plasma membrane and secretory proteins such as apolipoprotein B (*Apob*), apolipoprotein H (*Apoh*) and ceruloplasmin (*Cp*) (Figure 10A, red arrows). Inhibition of N-glycosylation modification of the glycoproteins results in their misfolding in the ER. Thus, the increase in binding of these proteins to BiP-FLAG in the livers of the Tm-treated BiP-FLAG mice provides direct evidence that misfolded ER proteins are co-precipitated with BiP-FLAG with anti-FLAG magnetic beads as components of BiP-interactome.

**Figure 10.**
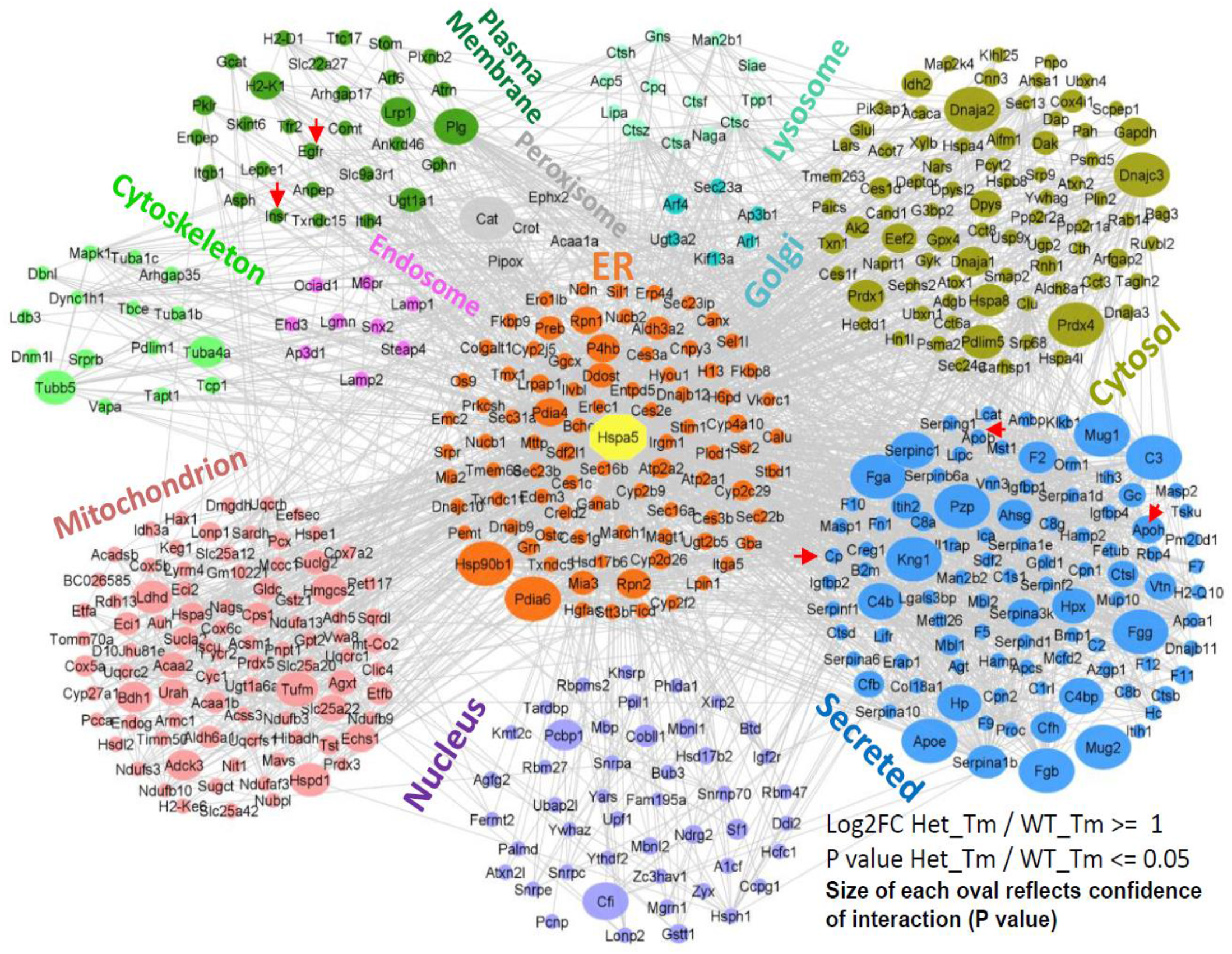
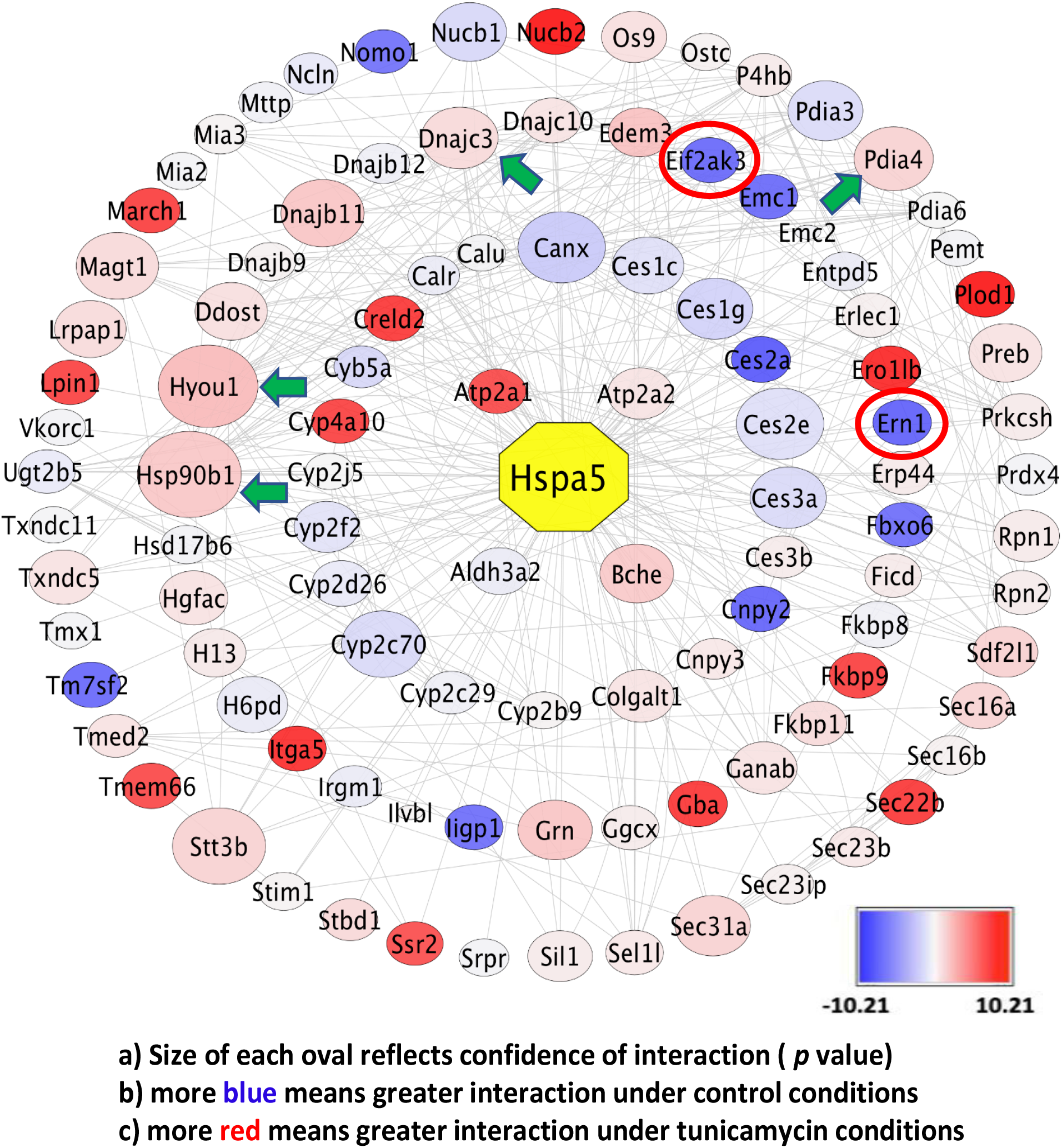
Proteomics analysis of BiP-FLAG complexes isolated from livers of vehicle- and tunicamycin (Tm)-treated BiP-FLAG heterozygous mice demonstrates the feasibility to use BiP-FLAG-expressing mice to isolate BiP-interactome and ER misfolded proteins. Mass Spec analysis was carried out using BiP-FLAG complexes isolated from the livers of vehicle-- and Tm-treated BiP-FLAG heterozygous and wild type mice described in Figure 9. **A**. A summary of proteins that exhibited augmented interaction with BiP-FLAG in response to Tm-treatment (red arrows, examples of glycoproteins); **B**. Effect of Tm-treatment on interactions of ER proteins with BiP-FLAG. Note, Tm treatment reduced BiP interactions with UPR sensors PERK (*Eif2ak3*) and IRE1α (*Ern1*) (red circles), indicating their release from BiP due to UPR activation, while promoting BiP interactions with chaperone proteins such as Grp94 (*Hsp90b1*), Grp170 (*Hyou1*), and P58^IPK^ (*Dnajc3*) (green arrows).

The ER proteins that displayed increased BiP-FLAG binding in response to TM-treatment are summarized in Figure 10B. Interestingly, the binding of PERK (*Eif2ak3*) and IRE1α (*Ern1*) to BiP-FLAG were greatly reduced after Tm-treatment (Figure 10 B, red circles). This is a significant finding because: a) For the first time, it provides *in vivo* evidence that these two major UPR transducers bind to BiP under non-stressed conditions and are released from BiP upon UPR activation; and b) It demonstrates the specificity of the proteins that co-IP with BiP-FLAG in BiP-FLAG-expressing mouse hepatocytes. As expected, the interactions of several ER chaperone proteins with BiP-FLAG, including Grp94 (*Hsp90b1*), GRP170 (*Hyou1*), and P58 (*Dnajc3*), were significantly increased in the Tm-treated BiP-FLAG Het livers, providing further evidence of the integrity of the BiP-interactome purified through the use of BiP-FLAG.

Together, the findings from AP-MS of the BiP-FLAG complexes isolated from livers of BiP-Flag mice under basal condition or Tm-induced ER stress demonstrate the feasibility of our unique *in vivo* model to detect changes in the BiP interaction network in response to ER stress and its usage to identify ER misfolded proteins.

### Pancreatic β cell specific BiP-FLAG mice reveal preferential binding of BiP-FLAG to proinsulin disulfide-linked complexes vs. monomeric proinsulin

In previous studies, we identified that proinsulin misfolding occurs even in healthy murine and human islets and increases further during prediabetes progression to Type 2 Diabetes (T2D) (19, 20). Proinsulin interacts with multiple ER chaperones and folding proteins in murine and human islets, among them, BiP was identified as a major binding partner of proinsulin in human healthy islets (12, 18). To identify the pathway for proinsulin folding *in vivo*, we generated pancreatic β cell specific BiP-Flag mice (*BiP-Flag;Ins1Cre*) by crossing whole body BiP-Flag knock-in mice with Ins1Cre mice, which constitutively express Cre recombinase from the *Ins1* gene (21).

First, we analyzed heterozygous *BiP-Flag;Ins1*^*Cre*^ (*BIP-Flag*^*fl*/+^;*Ins1*^*Cre*/+^) mice to check whether BiP-Flag expression in pancreas is β cell specific. We harvested pancreata from 8 wk-old Wt(+/+)*;Ins1*^*Cre*/+^ and *BIP-Flag*^*fl*/+^;*Ins1*^*Cre*/+^ male mice and performed IHC with anti-insulin antibody that recognizes both proinsulin and mature insulin, anti-Flag antibody and anti-PDIA6 antibody. We confirmed that BiP-Flag is only expressed in β cells and co-localized with PDIA6, an ER marker, indicating that BiP-FLAG is properly localized in the ER (Figure 11A–B).

**Figure 11.**
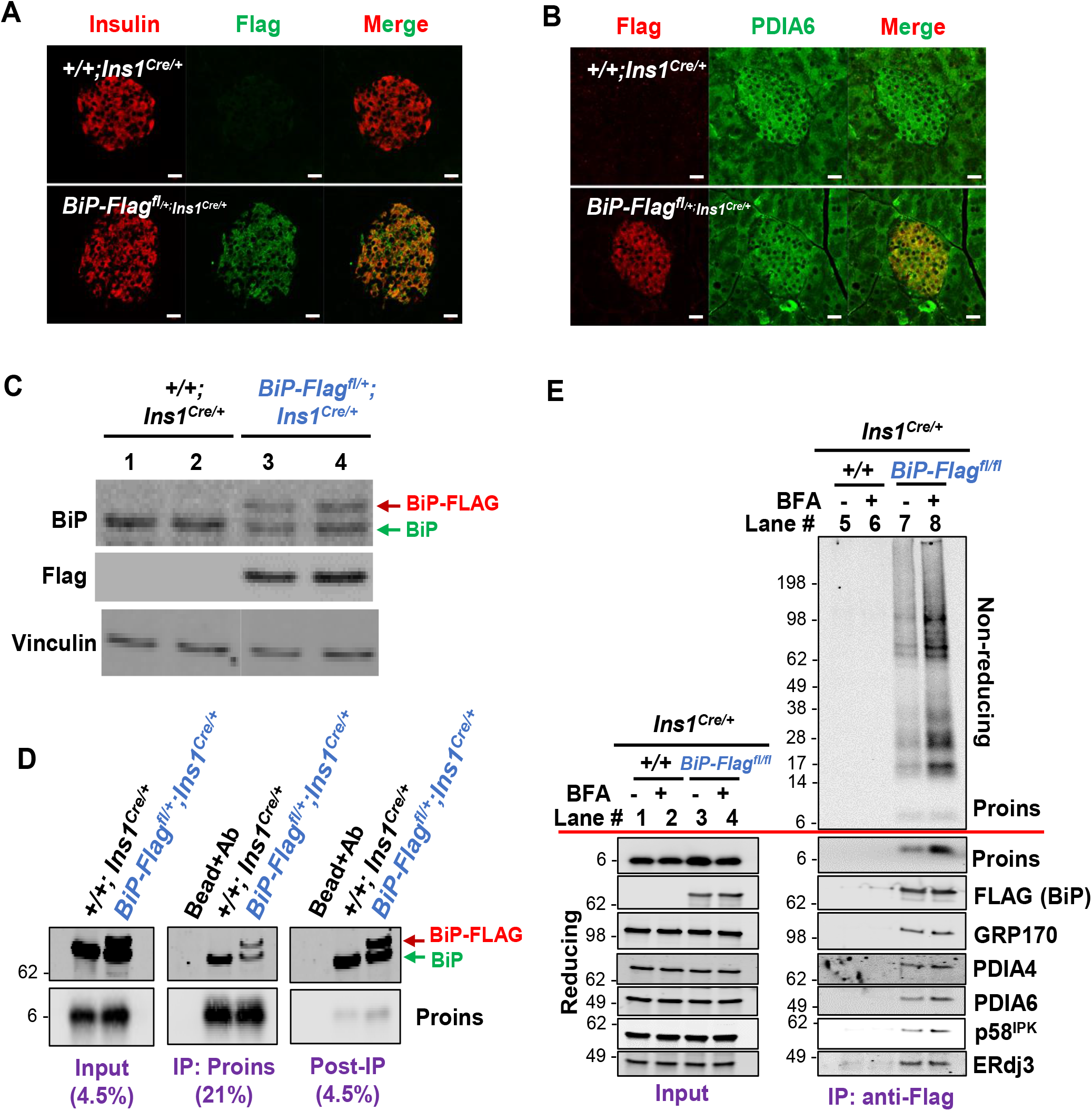
Demonstration of interaction of proinsulin with BiP-FLAG in β-cell specific BiP-Flag expressing mice. 1) BiP-Flag expressed specifically in β cells in *BiP-Flag*^*fl*/+^;*Ins1*^*Cre*/+^ mouse pancreas and co-localized with PDIA6 in the ER. Whole pancreas was isolated from 8 wk-old *Ins1*^*Cre*/+^ *male* mice with/without *BiP-Flag*^*fl*/+^ allele, fixed, and paraffin embedded. Then, sections were stained with Abs as indicated. **A.** Flag stained only in *BiP-Flag*^*fl*/+^;*Ins1*^*Cre*/+^ pancreas. Staining with Insulin Ab (recognizing both proinsulin and insulin) indicated that BiP-Flag is expressed in a β cell specific manner. **B.** Flag is co-stained with PDIA6, an ER marker in *BiP-Flag*^*fl*/+^;*Ins1*^*Cre*/+^ pancreas. Scale bar, 20μm. 2). BiP-Flag and BiP both bind to proinsulin in *BiP-Flag*^*fl*/+^;*Ins1*^*Cre*/+^ islets. **C.** Islets were isolated from 12 wk-old *Ins1*^*Cre*/+^ *male* mice with/without *BiP-Flag*^*fl*/+^ alleles. Lysates were analyzed by SDS-PAGE and immunoblotted with Abs indicated in the figure. **D.** Islets were isolated from 15 month-old *Ins1*^*Cre*^/+ *male* mice with/without *BiP-Flag*^*fl*/+^ alleles. Lysates were IP’d with proinsulin Ab and immunoblotted with Abs indicated. **E.** BiP binds HMW PI complexes in preference to monomeric PI and interacts with GRP170, PDIA4, PDIA6, p58^IPK^, and ERdj3, but not PDIA3. Islets were isolated from male +/+;*Ins1*^*Cre*/+^ or *BiP-Flag*^*fl/fl*^;*Ins1*^*Cre*/+^ mice after 10 days of HFD feeding. BFA (5μg/ml) was treated for 1h before harvesting islets. After IP with anti-Flag agarose beads, total lysates and IP elutes were analyzed by SDS-PAGE under non-reducing and reducing conditions with indicated Abs.

Next, we isolated islets from Wt(*+/+*)*;Ins1*^*Cre*/+^ and *BIP-Flag*^*fl*/+^;*Ins1*^*Cre*/+^ male mice to measure BiP and BiP-Flag protein expression and confirmed that BiP-Flag is only detected in *BIP-Flag*^*fl*/+^;*Ins1*^*Cre*/+^ islets and WT BiP and BiP-Flag protein were expressed at a comparable level (Figure 11C). Furthermore, we found both BiP and BiP-FLAG were co-IP’d with proinsulin from lysates of *BIP-Flag*^*fl*/+^;*Ins1*^*Cre*/+^ islets (Figure 11D), demonstrating that as expected, proinsulin interacts with both BiP and BiP-FLAG in a comparable manner. Thus, we generated homozygous *BiP-Flag;Ins1*^*Cre*^ mice to analyze BiP-FLAG/proinsulin proteome *in vivo* (Figure 11E).

We monitored body weight and blood glucose level of homozygous *BiP-Flag;Ins1*^*Cre*^ mice (*BIP-Flag*^*fl/fl*^;*Ins1*^*Cre*/+^) to determine whether BiP-Flag has any effect on mouse physiology. Expectedly, *BIP-Flag*^*fl/fl*^;*Ins1*^*Cre*/+^ mice did not show any significant differences in body weight and fed glucose level compared to *BIP-Flag*^*fl/fl*^ littermates. Furthermore, *BIP-Flag*^*fl/fl*^;*Ins1*^*Cre*/+^ did not show any abnormal sign in glucose tolerance tests as well as fasting blood glucose levels (Supplemental Figure 4).

We next performed FLAG IP to test if BiP-FLAG can pull down proinsulin as well as other ER chaperones which might be involved in proinsulin folding. To increase proinsulin synthesis *in vivo*, we fed *BIP-Flag*^*fl/fl*^;*Ins1*^*Cre*/+^ mice 45% HFD for 10 days before isolating islets. Isolated islets were incubated in complete media containing Brefeldin A (BFA) for 1h at 37°C to trap proinsulin in the ER. N-ethylmaleimide (NEM) was added into PBS for rinsing as well as the lysis buffer to alkylate free thiols. To halt BiP-Flag to substrate binding, 100 U/ml of hexokinase with 10 mM D-glucose was added to the lysis buffer. BiP-FLAG is only expressed in *BIP-Flag*^*fl/fl*^;*Ins1*^*Cre*/+^ islets (Figure 11E). We performed IP with goat-anti-Flag agarose beads to avoid of detecting mouse Ab chains and then analyzed IP elutes by SDS-PAGE under reducing and non-reducing conditions. As we expected, BFA treatment increased binding of BiP-FLAG to proinsulin under reducing gel conditions. Surprisingly, we noticed that BiP-FLAG preferentially bound to proinsulin disulfide-linked complexes vs. monomers as BFA treatment only increased proinsulin disulfide-linked complexes after IP and non-reducing gel SDS-PAGE (Figure 11E). Importantly, GRP170, PDIA4, PDIA6, p58^IPK^, and ERdj3 (but not PDIA3) were pulled down with proinsulin after BiP-Flag IP, indicating that these proteins might specifically assist proinsulin folding (Figure 11E). Further analysis will be necessary to identify whether any of these directly bind proinsulin and how they coordinate proinsulin folding with BiP.

## DISCUSSION

We demonstrated successful targeting of the endogenous *Hspa5* locus in mice. The phenotypic characterization identified no defect in hepatocyte function, ER function or BiP-FLAG localization to the ER. We believe that the epitope-tagged *Hspa5* locus will stimulate studies on BiP client specificity, function, and role in protein folding that was not previously possible. For example, FLAG IP of BiP-FLAG will permit answer to fundamental questions: **1)** Do different proteins misfold and bind BiP in healthy vs. diseased cells?; **2)** Do all inducers of ER stress generate similar protein misfolding consequences or are there differences depending on the degree or type of ER stress, or cell type?; **3)** What proteins exist in different complexes that contain BiP?; **4)** What are the kinetics of misfolded protein interaction and release from BiP?; and **5)** How do BiP interactions impact general ER processes? The BiP-FLAG mice will enable for the first-time delineation of folding pathways for any specific protein *in vivo.*

## ACKNOWLEDGEMENTS

R.J.K. is supported by NIH grants R01DK113171, R24DK110973, R37DK042394, R01CA198103, R01AG062190 and the SBP NCI Cancer Center Grant P30 CA030199. R.J.K. is a member of the UCSD DRC (P30 DK063491) and Adjunct Professor in the Department of Pharmacology, UCSD.

---

**Suppl. Figure 1.**
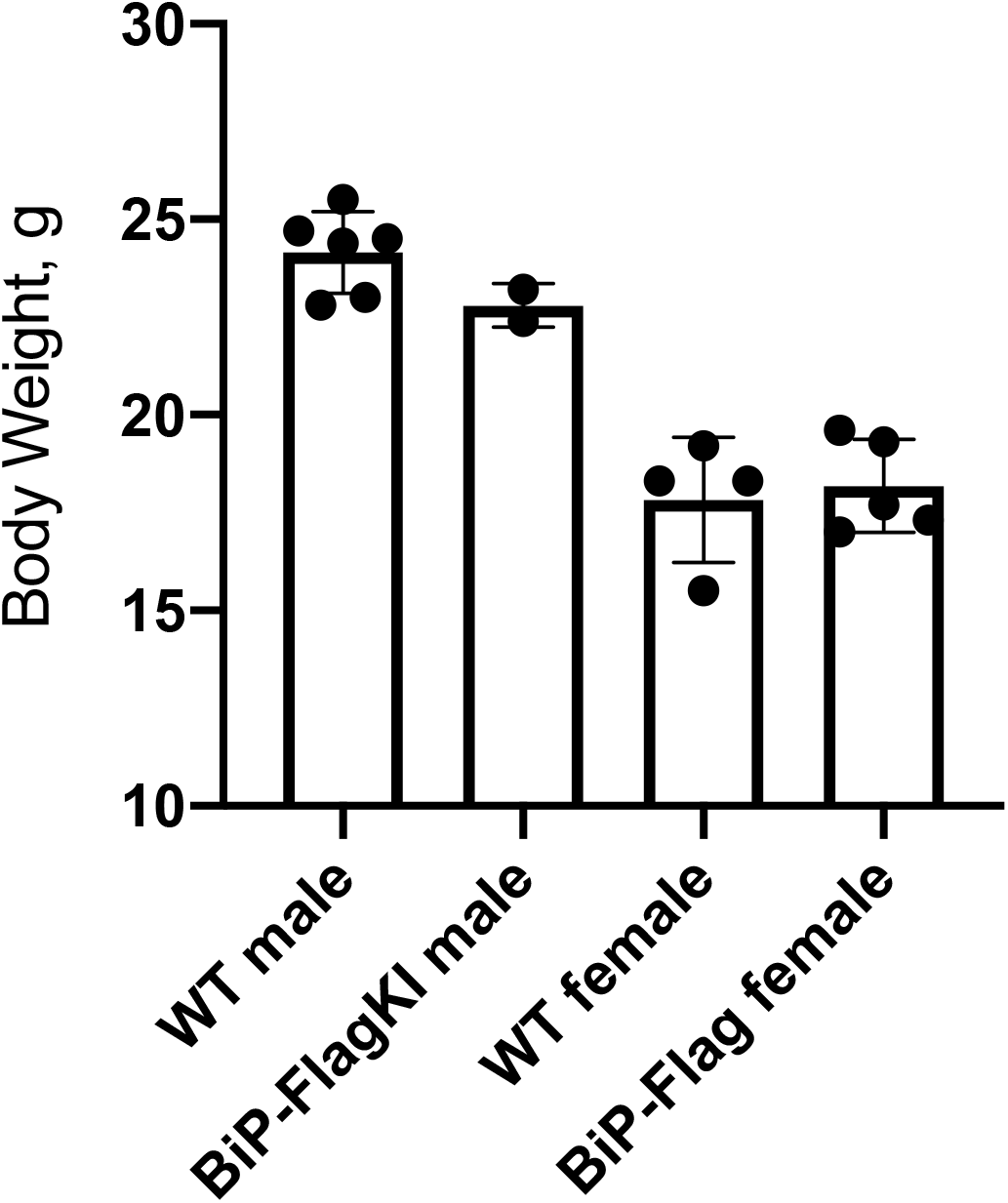
Body weight of WT and BiP-FLAG mice. Body weights were measured at 6-8 wks of age from WT and BiP-FLAG knock-in mice. WT, males=6, females=4. BiP-FLAG, males=2, females=5.

**Suppl. Figure 2.**
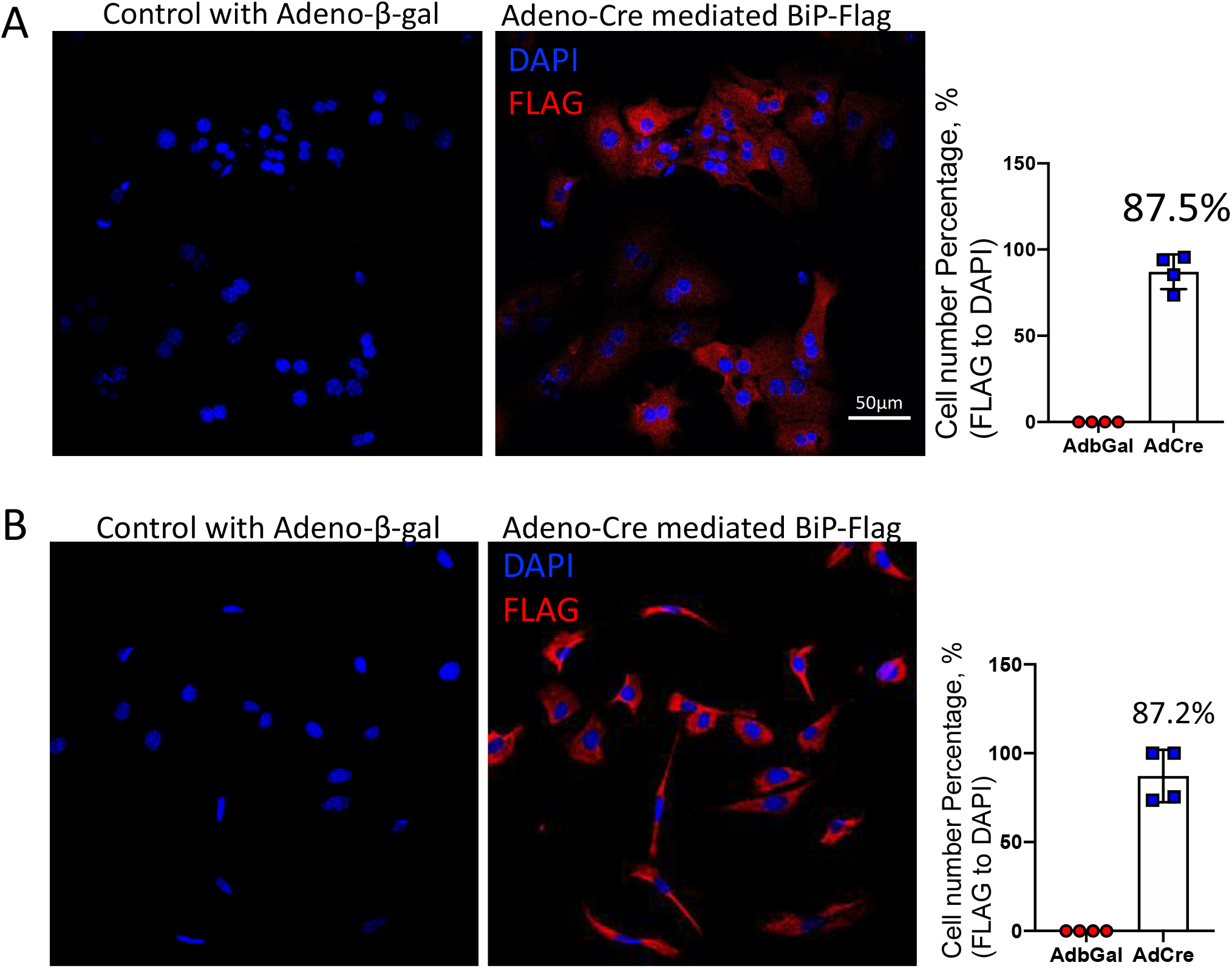
AAV infection efficiency in primary hepatocytes and fibroblasts. **A.** Hepatocytes were isolated from a 6-wk old female BiP-FLAG-Het mouse, plated onto 6 well plates and infected with the indicated adenoviruses at 4 h after plating. After 24 h, cells were fixed with formalin and stained with anti-FLAG antibody and DAPI. **B.** Fibroblasts were isolated from a 6-wk old female BiP-FLAG-Het mouse, plated onto 6 well plates and infected with the indicated adenoviruses at 7 days after plating. At 24 h after Ad-infection, cells were fixed with formalin and staining with anti-FLAG antibody and DAPI. Four Images were randomly captured from each group by Zeiss 710 confocal microscopy. Scale bar, 50 μm. Quantification was performed by Image J. Quantification of the ratio of FLAG-positive cells to DAPI positive cells is shown in each graph (right).

**Suppl. Figure 3.**
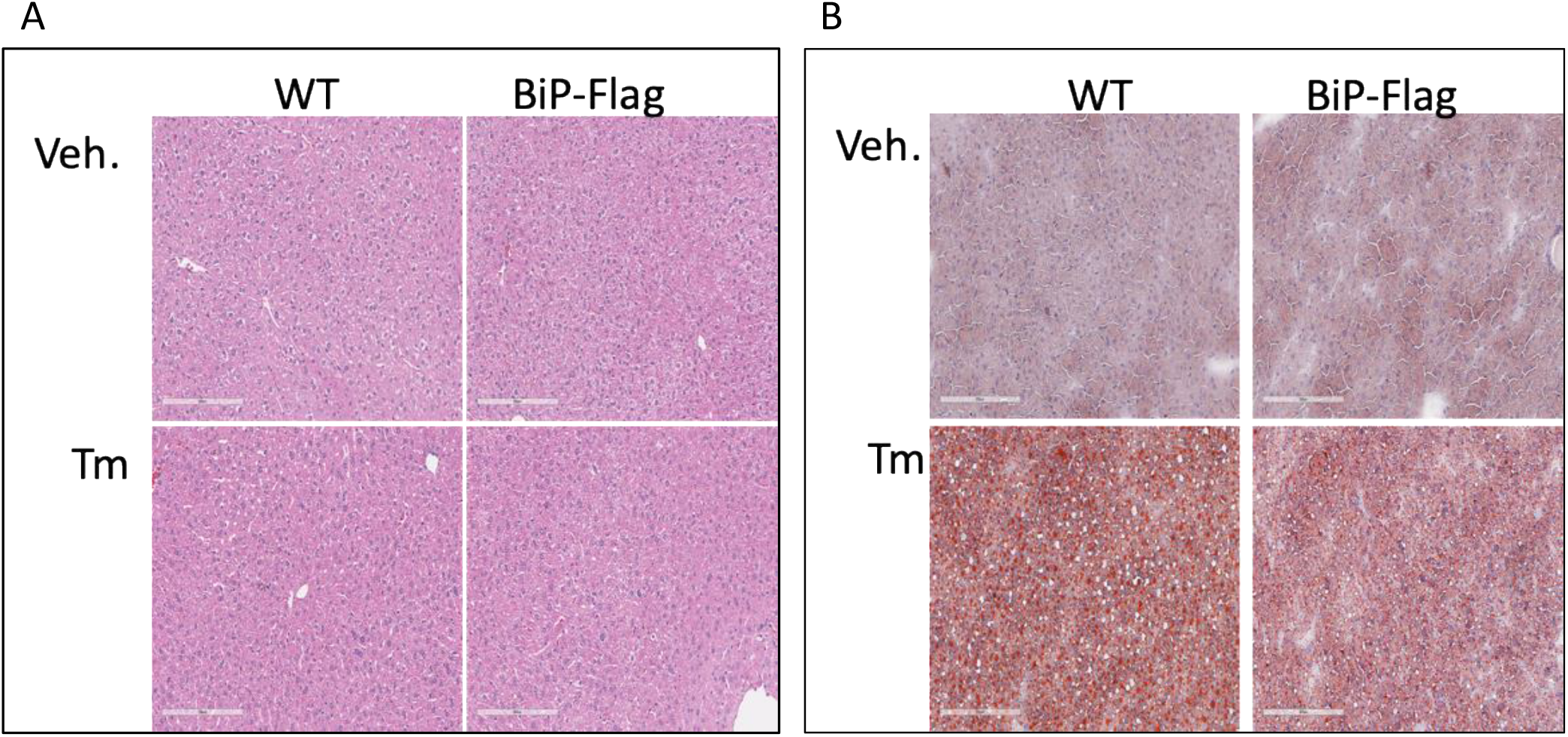
Liver histology of AAV-TBG-Cre infected BiP-FLAG-Het mice. **A.** Morphology of hepatocytes based on H&E stained liver sections of experimental mice. **B.** Pathohistological analysis and morphology of hepatocytes based on Oil Red O stained liver sections of experimental mice. Experiments were performed with 2 mice in each group. Stained sections were scanned by Aperio, Leica Biosystems. The scale bar is 200 μm.

**Suppl Figure 4.**
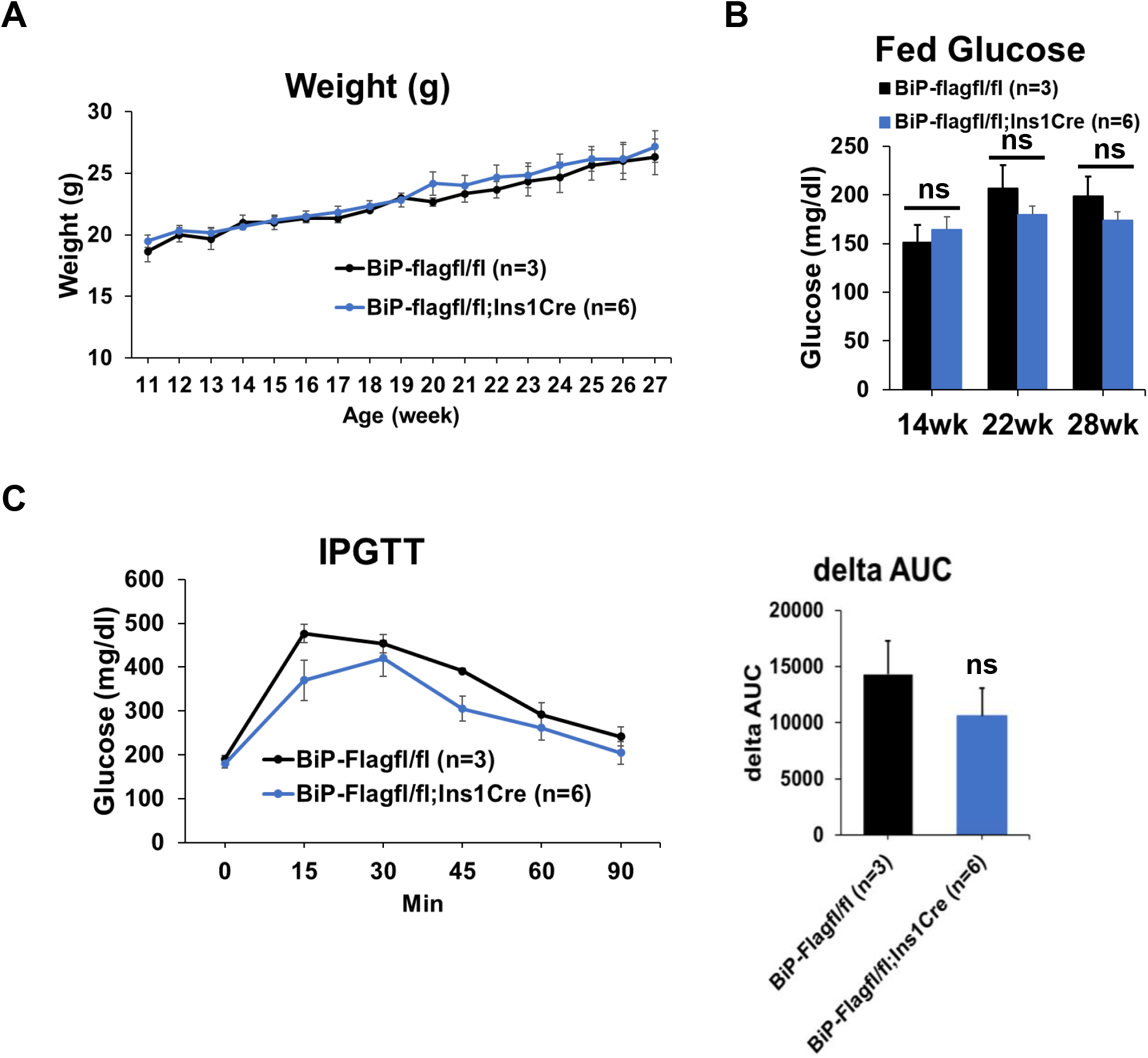
No significant difference is observed in body weight, blood glucose level, and glucose tolerance in *BiP-Flag/BiP-Flag;Ins1*^*Cre*^/+ mice compared to *BiP-Flag/BiP-Flag* mice. **A.** Body weight (g) **B.** Fed blood glucose level (mg/dL) **C.** Glucose tolerance test at 11 wks. After 4h fasting, blood glucose levels (mg/dL) were measured by tail-bleeding before and after glucose injection. Mean ± SEM. ns, not significant.

**Suppl. Table 1.**
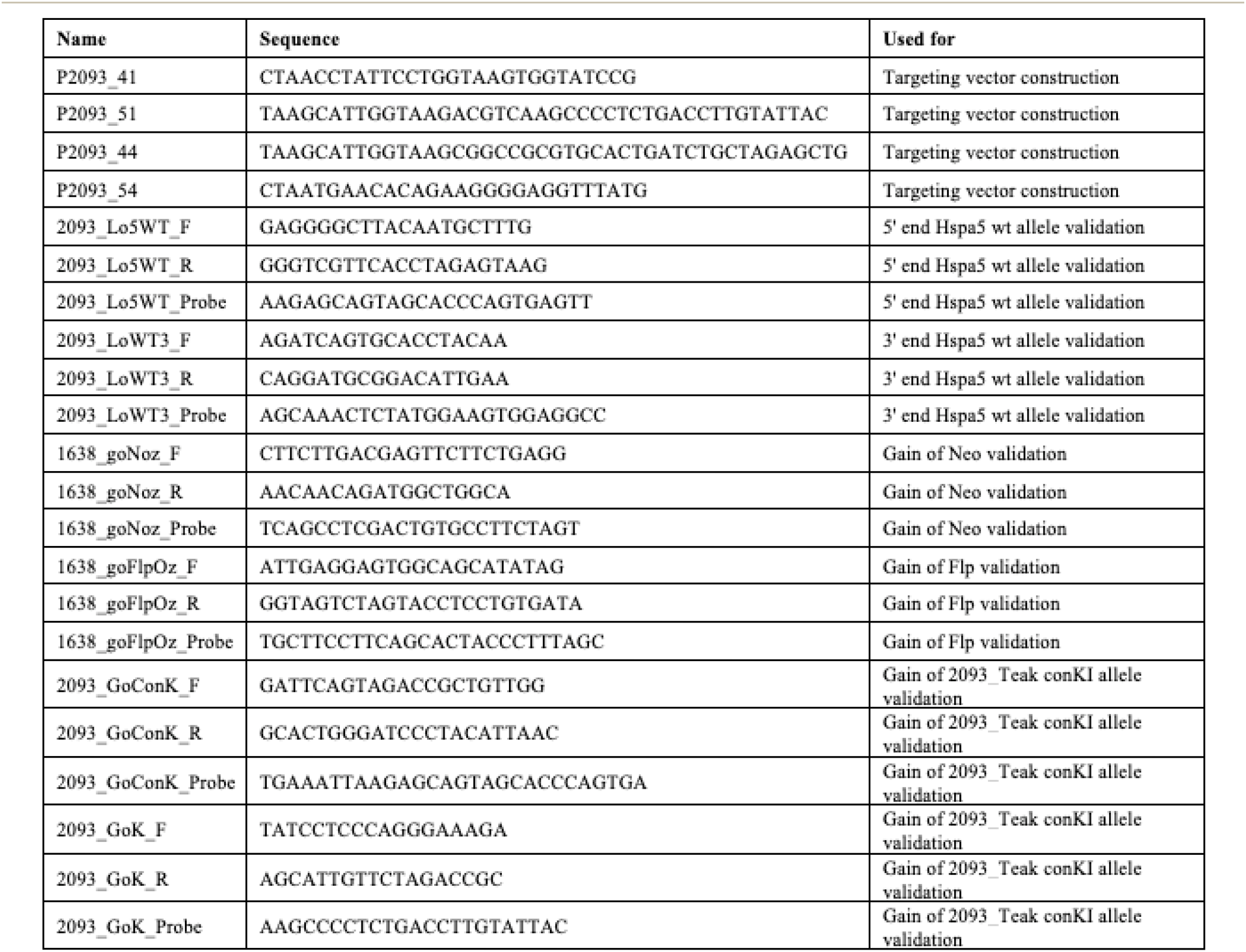
Primers used.

## REFERENCES

1. Hebert DN, Molinari M. In and out of the ER: protein folding, quality control, degradation, and related human diseases. Physiol Rev. 2007;87(4):1377–408. Epub 2007/10/12. doi: 10.1152/physrev.00050.2006. PubMed PMID: 17928587.

2. Kaufman RJ. Orchestrating the unfolded protein response in health and disease. J Clin Invest. 2002;110(10):1389–98. Epub 2002/11/20. doi: 10.1172/JCI16886. PubMed PMID: 12438434; PMCID: PMC151822.

3. Ron D, Walter P. Signal integration in the endoplasmic reticulum unfolded protein response. Nat Rev Mol Cell Biol. 2007;8(7):519–29. Epub 2007/06/15. doi: 10.1038/nrm2199. PubMed PMID: 17565364.

4. Haas IG, Wabl M. Immunoglobulin heavy chain binding protein. Nature. 1983;306(5941):387–9. Epub 1983/11/24. doi: 10.1038/306387a0. PubMed PMID: 6417546.

5. Munro S, Pelham HR. An Hsp70-like protein in the ER: identity with the 78 kd glucose-regulated protein and immunoglobulin heavy chain binding protein. Cell. 1986;46(2):291–300. Epub 1986/07/18. doi: 10.1016/0092-8674(86)90746-4. PubMed PMID: 3087629.

6. Bertolotti A, Zhang Y, Hendershot LM, Harding HP, Ron D. Dynamic interaction of BiP and ER stress transducers in the unfolded-protein response. Nat Cell Biol. 2000;2(6):326–32. Epub 2000/06/15. doi: 10.1038/35014014. PubMed PMID: 10854322.

7. Dorner AJ, Bole DG, Kaufman RJ. The relationship of N-linked glycosylation and heavy chain-binding protein association with the secretion of glycoproteins. J Cell Biol. 1987;105(6 Pt 1):2665–74. Epub 1987/12/01. doi: 10.1083/jcb.105.6.2665. PubMed PMID: 3121636; PMCID: PMC2114744.

8. Kozutsumi Y, Segal M, Normington K, Gething MJ, Sambrook J. The presence of malfolded proteins in the endoplasmic reticulum signals the induction of glucose-regulated proteins. Nature. 1988;332(6163):462–4. Epub 1988/03/31. doi: 10.1038/332462a0. PubMed PMID: 3352747.

9. Dorner AJ, Wasley LC, Kaufman RJ. Increased synthesis of secreted proteins induces expression of glucose-regulated proteins in butyrate-treated Chinese hamster ovary cells. J Biol Chem. 1989;264(34):20602–7. Epub 1989/12/05. PubMed PMID: 2511206.

10. Dorner AJ, Wasley LC, Kaufman RJ. Overexpression of GRP78 mitigates stress induction of glucose regulated proteins and blocks secretion of selective proteins in Chinese hamster ovary cells. EMBO J. 1992;11(4):1563–71. Epub 1992/04/01. PubMed PMID: 1373378; PMCID: PMC556605.

11. Ng DT, Watowich SS, Lamb RA. Analysis in vivo of GRP78-BiP/substrate interactions and their role in induction of the GRP78-BiP gene. Mol Biol Cell. 1992;3(2):143–55. Epub 1992/02/01. doi: 10.1091/mbc.3.2.143. PubMed PMID: 1550958; PMCID: PMC275514.

12. Scheuner D, Vander Mierde D, Song B, Flamez D, Creemers JW, Tsukamoto K, Ribick M, Schuit FC, Kaufman RJ. Control of mRNA translation preserves endoplasmic reticulum function in beta cells and maintains glucose homeostasis. Nat Med. 2005;11(7):757–64. Epub 2005/06/28. doi: 10.1038/nm1259. PubMed PMID: 15980866.

13. Hidvegi T, Schmidt BZ, Hale P, Perlmutter DH. Accumulation of mutant alpha1-antitrypsin Z in the endoplasmic reticulum activates caspases-4 and −12, NFkappaB, and BAP31 but not the unfolded protein response. J Biol Chem. 2005;280(47):39002–15. Epub 2005/09/27. doi: 10.1074/jbc.M508652200. PubMed PMID: 16183649.

14. Kontgen F, Suss G, Stewart C, Steinmetz M, Bluethmann H. Targeted disruption of the MHC class II Aa gene in C57BL/6 mice. Int Immunol. 1993;5(8):957–64. Epub 1993/08/01. doi: 10.1093/intimm/5.8.957. PubMed PMID: 8398989.

15. Koentgen F, Lin J, Katidou M, Chang I, Khan M, Watts J, Mombaerts P. Exclusive transmission of the embryonic stem cell-derived genome through the mouse germline. Genesis. 2016;54(6):326–33. Epub 2016/03/26. doi: 10.1002/dvg.22938. PubMed PMID: 27012318; PMCID: PMC5084746.

16. Zhou H, Zeng Z, Koentgen F, Khan M, Mombaerts P. The testicular soma of Tsc22d3 knockout mice supports spermatogenesis and germline transmission from spermatogonial stem cell lines upon transplantation. Genesis. 2019;57(6):e23295. Epub 2019/04/20. doi: 10.1002/dvg.23295. PubMed PMID: 31001916; PMCID: PMC6617806.

17. Wang S, Chen Z, Lam V, Han J, Hassler J, Finck BN, Davidson NO, Kaufman RJ. IRE1alpha-XBP1s induces PDI expression to increase MTP activity for hepatic VLDL assembly and lipid homeostasis. Cell Metab. 2012;16(4):473–86. Epub 2012/10/09. doi: 10.1016/j.cmet.2012.09.003. PubMed PMID: 23040069; PMCID: PMC3569089.

18. Tran DT, Pottekat A, Mir SA, Loguercio S, Jang I, Campos AR, Scully KM, Lahmy R, Liu M, Arvan P, Balch WE, Kaufman RJ, Itkin-Ansari P. Unbiased Profiling of the Human Proinsulin Biosynthetic Interaction Network Reveals a Role for Peroxiredoxin 4 in Proinsulin Folding. Diabetes. 2020;69(8):1723–34. Epub 2020/05/28. doi: 10.2337/db20-0245. PubMed PMID: 32457219; PMCID: PMC7372081.

19. Jang I, Pottekat A, Poothong J, Yong J, Lagunas-Acosta J, Charbono A, Chen Z, Scheuner DL, Liu M, Itkin-Ansari P, Arvan P, Kaufman RJ. PDIA1/P4HB is required for efficient proinsulin maturation and ss cell health in response to diet induced obesity. Elife. 2019;8. Epub 2019/06/12. doi: 10.7554/eLife.44528. PubMed PMID: 31184304; PMCID: PMC6559792.

20. Arunagiri A, Haataja L, Pottekat A, Pamenan F, Kim S, Zeltser LM, Paton AW, Paton JC, Tsai B, Itkin-Ansari P, Kaufman RJ, Liu M, Arvan P. Proinsulin misfolding is an early event in the progression to type 2 diabetes. Elife. 2019;8. Epub 2019/06/12. doi: 10.7554/eLife.44532. PubMed PMID: 31184302; PMCID: PMC6559786.

21. Thorens B, Tarussio D, Maestro MA, Rovira M, Heikkila E, Ferrer J. Ins1(Cre) knock-in mice for beta cell-specific gene recombination. Diabetologia. 2015;58(3):558–65. Epub 2014/12/17. doi: 10.1007/s00125-014-3468-5. PubMed PMID: 25500700; PMCID: PMC4320308.

